# Shedding Light on the Dark Genome: Insights into the Genetic, CRISPR-based, and Pharmacological Dependencies of Human Cancers and Disease Aggressiveness

**DOI:** 10.1101/2023.08.15.552589

**Authors:** Doris Kafita, Panji Nkhoma, Kevin Dzobo, Musalula Sinkala

**Affiliations:** University of Zambia, School of Health Sciences, Department of Biomedical Sciences, Lusaka, Zambia; Wound and Keloid Scarring Research Unit, Hair and Skin Research Laboratory, Division of Dermatology, Department of Medicine, The South African Medical Research Council, University of Cape Town, Cape Town, South Africa; University of Cape Town, Faculty of Health Sciences, Institute of Infectious Disease and Molecular Medicine and Department of Integrative Biomedical Sciences, Computational Biology Division, Cape Town, South Africa

## Abstract

Investigating the human genome is vital for identifying risk factors and devising effective therapies to combat genetic disorders and cancer. Despite the extensive knowledge of the “light genome”, the poorly understood “dark genome” remains understudied. In this study, we integrated data from 20,412 protein-coding genes in Pharos and 8,395 patient-derived tumours from The Cancer Genome Atlas (TCGA) to examine the genetic and pharmacological dependencies in human cancers and their treatment implications. We discovered that dark genes exhibited high mutation rates in certain cancers, similar to light genes. By combining the drug response profiles of cancer cells with cell fitness post-CRISPR-mediated gene knockout, we identified the crucial vulnerabilities associated with both dark and light genes. Our analysis also revealed that tumours harbouring dark gene mutations displayed worse overall and disease-free survival rates than those without such mutations. Furthermore, dark gene expression levels significantly influenced patient survival outcomes. Our findings demonstrated a similar distribution of genetic and pharmacological dependencies across the light and dark genomes, suggesting that targeting the dark genome holds promise for cancer treatment. This study underscores the need for ongoing research on the dark genome to better comprehend the underlying mechanisms of cancer and develop more effective therapies.

## Introduction

The human genome comprises approximately 19,000-25,000 protein-coding gene sequences, accounting for 1-2% of the genome^1–5^. Perturbations within the genome can result in changes in cellular phenotypes and behavioural alterations^6–8^. Such disruptions in protein-coding regions are closely associated with the onset of and susceptibility to human diseases, including cancer and other genetic disorders^9,10^. For example, tumour development and progression are strongly influenced by activated oncogenes and the inactivation of tumour suppressor genes, thereby providing cancer-specific hallmarks^11–16^. The introduction of high-throughput next-generation sequencing techniques has led to the identification of numerous known and novel causal genes^17,18^ through the rapid sequencing of complete genomes and generation of extensive genome data^19–22^. Despite the increasing number of systematic studies on cancer and genetic disorders, cancer remains the second leading cause of death worldwide^23–26^.

The “dark genome”, “ignorome”, or “Tdark” comprises protein-coding genes with limited or no known function in literature, representing over a third of all genes^27^. This vast amount of unexplored genetic information highlights the significant gaps in our understanding of their roles and importance^27–29^. The lack of knowledge surrounding these genes poses a considerable obstacle to the advancement of personalised medicine, despite the ever-growing comprehension of the human genome^28,30^. While our understanding of the human genome has undeniably facilitated the diagnosis of genetic diseases and the development of targeted therapies for various pathological disorders, such as cancer^5^, the dark genome represents a substantial challenge to the progress of personalised medicine^27^. The inability to decipher the functions and significance of these genes limits our ability to tailor medical treatments based on individual genetic variations, thereby impeding the full realisation of personalised medicine’s potential^27^. Overcoming this obstacle requires extensive research aimed at unravelling the mysteries of the dark genome, shedding light on its function, and unlocking its therapeutic potential.

Large molecular profiling projects, such as The Cancer Genome Atlas (TCGA) project^31^, have extensively profiled human cancers, providing valuable insights into their molecular landscapes and potential therapeutic targets. The Achilles project utilises CRISPR technology to explore gene dependencies in cancer cells^32^, whereas the Genomics of Drug Sensitivity in Cancer (GDSC)^33^ and Cancer Cell Line Encyclopaedia (CCLE)^34^ projects screen thousands of cancer cell lines for small-molecule inhibitor responses. These efforts have generated vast publicly accessible data for advancing cancer understanding and treatment and uncovering the mysteries of the dark genome. The Illuminating the Druggable Genome (IDG) project has compiled dark gene data from over 60 sources to identify new therapeutic targets and advance cancer understanding^29,35^. This effort led to the development of the Target Central Resource Database (TCRD), which is a comprehensive database that integrates various data types. To facilitate easy exploration and sharing of this data, a web-based platform called Pharos was created^36^.

Integrating the different datasets from TCGA, Achilles, GDSC, CCLE, and IDG projects is essential for illuminating the dark genome and overcoming obstacles posed by unstudied human genes, thus facilitating the advancement of personalised medicine. In this study, we extensively analysed the distribution of genes in both light and dark genomes, investigated the mutation frequency of dark genes across various human cancer types, and assessed their impact on the chemosensitivity of cancer cell lines and aggressiveness of cancer. Our comprehensive analysis aimed to provide valuable insights into the role of the dark genome in human cancers and contribute to a better understanding of the genetic landscape of these diseases. Furthermore, by leveraging information from large-scale molecular profiling projects, we underscore the importance of continued research on the dark genome and the integration of multiple datasets to enhance our understanding of its impact on cancer, ultimately promoting more effective, targeted therapies for cancer and other genetic disorders.

## Results

### Distribution and potential of dark genes as therapeutic targets

We obtained human gene classification information based on target development levels (TDLs) from Pharos [version 3.15.1] (https://pharos.nih.gov), a multimodal web interface that presents data from the Target Central Resource Database (TCRD)^29,36^. Pharos classified genes/proteins into four TDLs: light genes (Tbio [n = 12,058], Tclin [n = 704], and Tchem [n = 1,971]), and dark genes (Tdark [n = 5,679]) (see Supplementary Figure 1a and “Methods” section for the description of TDLs). Our evaluation of TDL genes revealed that only 3.5% are currently utilised as drug targets, suggesting that many genes have the potential to be developed as drug targets (Tchem), and that a significant proportion (27.8%) of the dark genome encoding Tdark proteins remains to be understood. This study further filtered the dataset to include genes with information on publications, monoclonal antibodies, and antibody counts. The final datasets encompassed 19,387 genes, including light genes (Tbio [n = 11,724], Tclin [n = 689], and Tchem [n = 1,946]) and dark genes (Tdark [n = 5,028]). The distribution of genes within each development level is illustrated in Figure 1a (see Supplementary File 1).

**Figure 1.**
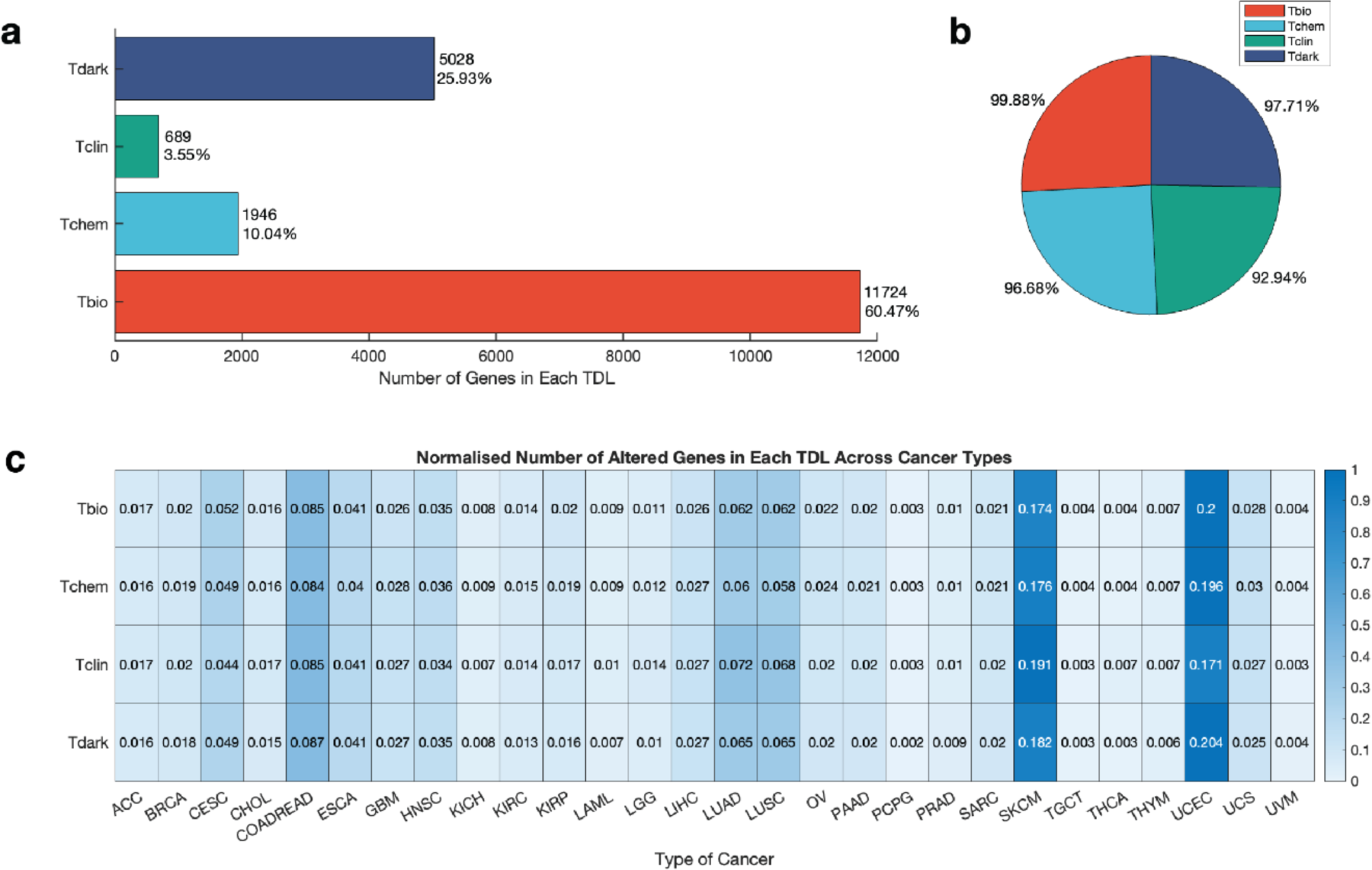
Distribution of dark and light genes and their mutations in human cancers. **a.** Number of genes in each target development level. **b.** Percentage of cancers with mutations at each target developmental level. **c.** Normalized number of mutated genes at each target development level across the 28 cancer types. ACC: Adenoid cystic carcinoma; BRCA: Breast cancer; CESC: Cervical squamous cell carcinoma; CHOL: Cholangiocarcinoma; COADREAD: Colorectal cancer; ESCA: Oesophageal carcinoma; GBM: Glioblastoma multiforme; HNSC: Head and neck squamous cell carcinoma; KICH: Kidney chromophobe; KIRC: Kidney renal clear cell carcinoma; KIRP: Kidney renal papillary cell carcinoma; LAML: Acute myeloid leukaemia; LGG: Brain lower grade glioma; LIHC: Liver hepatocellular carcinoma; LUAD: Lung adenocarcinoma; LUSC: Lung squamous cell carcinoma; OV: Ovarian serous cystadenocarcinoma; PAAD: Pancreatic adenocarcinoma; PCPG: Pheochromocytoma and paraganglioma; PRAD: Prostate adenocarcinoma; SARC: Sarcoma; SKCM: Skin cutaneous melanoma; TGCT: Testicular germ cell tumours; THCA: Thyroid carcinoma; THYM: Thymoma; UCEC: Uterine corpus endometrial carcinoma; UCS: Uterine carcinosarcoma; UVM: Uveal melanoma.

We observed a reduction in dark genes between TCRD versions 4.3.4 and 6.13.4, with version 4.3.4 containing 7,003 Tdark genes and version 6.13.4 with 5,679 Tdark genes, indicating a decrease of 1,324 genes (see Supplementary Figure 1b). Most genes, including *ACTR8*, *CSMD3*, *LSM3*, and *AASDH*, which were previously classified as Tdark, are now categorised as Tbio, likely because of new information regarding their functions. This reduction in dark gene expression between the two database versions reflects the progress in our understanding of the human genome and its potential as a source of novel therapeutic targets. Furthermore, this reveals that what is called “dark genes” currently may have a function after careful analysis and further studies.

### Research focus and trends in target development levels

To determine the research focus for each development level, we performed a one-way analysis of variance (ANOVA) to compare the mean scores of target development levels on publication count, antibody count, and monoclonal antibody count. The statistical analysis revealed significant main effects of target development levels on publication count (F(3, 19383) = 6.07 x 10^3^, p = 1 × 10^−300^) (Figure 2a), antibody count (F(3, 19383) = 3.85 x 10^3^, p = 1 × 10^−300^) (Figure 2b), and monoclonal antibody count (F(3,19383) = 2.62 x10^3^, p = 1 × 10^−300^) (Figure 2c). These results suggest significant impacts of target development levels on publication, antibody, and monoclonal antibody counts. We further analysed the differences between target development levels using Bonferroni post hoc comparisons, revealing that Tclin had the highest publication count (mean = 4.53) and Tdark the lowest (mean = 1.84). Similarly, Tclin had the highest antibody count (mean = 5.40), and Tdark had the lowest (mean = 2.85). Tclin also had the highest monoclonal antibody count (mean = 3.21), while Tdark had the lowest (mean = 0.39). Our findings suggest that the research focus is consistently high in Tclin, followed by Tchem, Tbio, and Tdark, offering valuable insights into research priorities and trends within different development levels and informing future research efforts.

**Figure 2.**
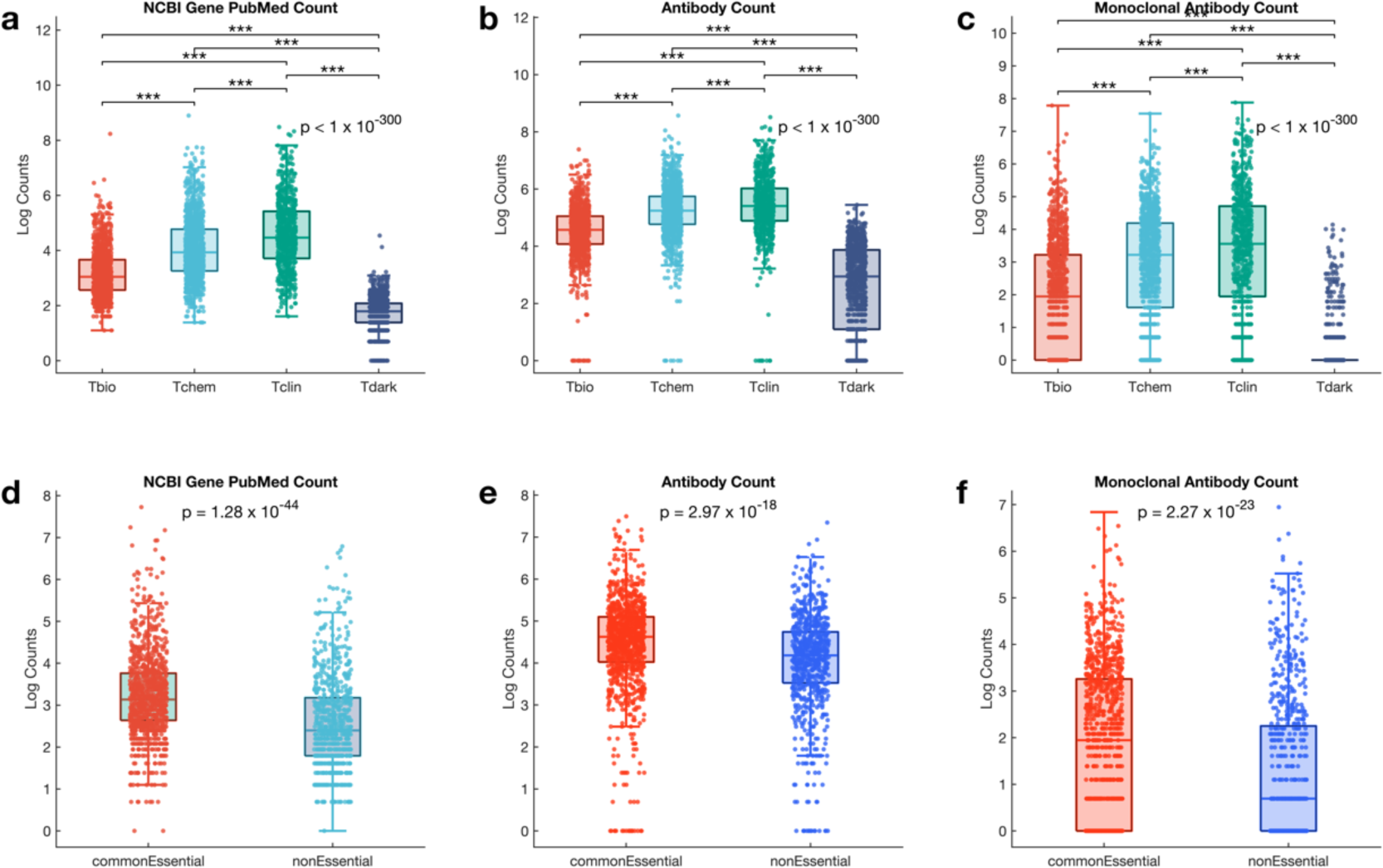
Research distribution across target development levels. Comparison of the **a.** publication count, **b.** antibody count, and **c.** Monoclonal antibody count among the four target development levels. Comparison of the **d.** publication count, **e.** antibody count and **f.** Monoclonal antibody counts between common essential (n = 2073) and non-essential (n = 722) genes. The boxplots indicate the distribution of the publication, antibody, and monoclonal antibody counts. On each box, the central mark indicates the median, and the bottom edge represents the 25th percentile, whereas the top edge of the box represents the 75th percentile. The whiskers extend to the most extreme data points that are not considered outliers. The scatter points within each box plot show the overall distribution of data points.

To identify potential differences between essential and non-essential genes in research and development efforts, we assessed the mean publication count, antibody count, and monoclonal antibody count between the common essential (N = 2073) and non-essential (N = 722) genes in the Achilles project dataset. We found significantly higher mean publication counts for common essential genes (mean = 3.25) compared to non-essential genes (mean = 2.56) (Welch test: t = 14.66, p = 1.28 × 10^−44^) (Figure 2d). Additionally, both antibody and monoclonal antibody counts were significantly higher in common essential genes (mean antibody count = 4.47, mean monoclonal antibody count = 1.91) than in non-essential genes (mean antibody count = 4.02, mean monoclonal antibody count = 1.22), antibody count (t = 8.86, p = 2.97 × 10^−18^), and monoclonal antibody count (t = 14.66, p = 2.27 × 10^−23^) (Figure 2e and 2f), suggesting that essential genes receive more research attention.

### Dark genes are as frequently mutated across human cancers as light genes

To investigate the extent of mutations in dark and light genes in cancer, we obtained a dataset of 8,395 human cancer cases across 28 primary tumours from TCGA^31^, consisting of gene copy number alterations and somatic mutations. We integrated this information with the TDL classification of genes from Pharos and assessed the number of cancers with mutations in each TDL class. Our results revealed that most cancers harboured Tbio gene mutations (99.88%), followed by Tdark (97.71%), Tchem (96.68%), and Tclin (92.94%) (Figure 1b and Supplementary File 1).

We further compared the number of mutated dark genes to light genes across each of the 28 cancer types. Our findings showed that the extent of dark gene mutations varied greatly depending on cancer type. Specifically, UCEC (Uterine corpus endometrial carcinoma) exhibited the highest number of dark gene mutations (20.39%), followed by SKCM (Skin cutaneous melanoma) (18.25%), COADREAD (Colorectal cancer) (8.66%), LUAD (Lung adenocarcinoma) (6.52%), LUSC (Lung squamous cell carcinoma) (6.52%), and CESC (Cervical squamous cell carcinoma) (4.90%). A more granular analysis of SKCM revealed the following normalised and absolute values: 0.17 (624) Tbio genes, 0.18 (118) Tchem genes, 0.19 (56) Tclin genes, and 0.18 (179) Tdark genes were highly mutated. Likewise, for UCEC, we found the following normalised and absolute values: 0.20 (716) Tbio genes, 0.20 (131) Tchem genes, 0.17 (50) Tclin genes, and 0.20 (200) Tdark genes were highly mutated (Figure 1c, also see Supplementary Figure 1c). Our findings suggest numerous potential therapeutic targets for treating these cancer types, particularly Tbio, Tchem and Tdark genes.

Furthermore, our analysis delved into the specific mutated dark genes within each cancer type, shedding light on the most frequently affected genes. In SKCM, the following Tdark genes exhibited the highest mutation rates: *PKHD1L1* (52.07%), *DNAH9* (45.18%), *THSD7B* (44.90%), *DNAH3* (43.52%), and *RP1* (42.98%). Similarly, in UCEC, the five most frequently mutated Tdark genes, were *MDN1*(20.71%), *DNHD1* (19.72%), *DNAH3* (19.53%), *SSPO* (19.33%), and *DNAH9* (19.13%). Within COADREAD, the predominant mutations were observed in *DCHS2* (16.98%), *DNAH17* (13.74%), *MDN1* (13.36%), *UNC13C* (13.36%), and *SSPO* (13.17%). For LUAD, the frequently mutated Tdark genes included *DNAH9* (22.07%), *SSPO* (18.49%), *BAGE2* (18.09%), *ZNF831* (16.90%), and *PKHD1L1* (15.71%). In the case of LUSC, the most mutated Tdark genes were *PKHD1L1* (23.82%), *DNAH9* (19.74%), *BAGE2* (18.24%), *ZNF804B* (17.38%), and *SPHKAP* (17.38%). Similarly, CESC exhibited frequent mutations in *MDN1* (8.36%), *DNAH3* (8.36%), *DNAH6* (8.36%), *SSPO* (8.00%), and *FSIP2* (7.63%) among Tdark genes (Supplementary File 1).

Notably, *PKHD1L1* emerged as the most frequently mutated Tdark gene across all 28 cancer types, highlighting its potential significance in cancer development and progression. Conversely, *TP53* (Tchem) stood out as the most frequently mutated light gene across all 28 cancer types, especially in OV (Ovarian cancer) (94.5%) (Supplementary File 1). This finding aligns with previous observations and reinforces the established association between *TP53* mutations and OV^37–45^.

### Dark genes strongly impact the fitness of cancer cells

To assess the importance of dark genes compared with light genes, we used Achilles^32^ data to analyse the impact of genes in each target development level (Tclin, Tchem, Tbio, and Tdark) on cancer cell lines. In the Achilles project, gene dependency scores measure the effect of gene perturbation on cell fitness, where a score close to −1 indicates reduced fitness (increased dependency), a score close to 1 indicates increased fitness (reduced dependency), and a score of 0 indicates no change in fitness (independence).

A one-way ANOVA test was performed to compare the mean gene dependency scores among the target development levels, revealing a statistically significant difference, F(3, 14120423) = 5.29 x 10^4^, P < 1 x 10^-300^. The mean gene dependency scores for Tclin, Tchem, Tbio, and Tdark were −0.10, −0.14, −0.17, and −0.07, respectively. Tbio had the lowest mean gene dependency score, signifying the greatest negative impact on cell viability, followed by Tchem, Tclin, and Tdark, with the least negative impact. However, the effect sizes suggest that this difference accounted for a relatively small proportion of the overall variance. Specifically, the eta-squared (η^2^) value was 0.0111, indicating that 1.11% of the total variance in our outcome variable can be attributed to group membership. The omega-squared (ω^2^) value was 0.0111, suggesting a similar proportion of explained variance when adjusted for bias. Given the very large sample size of 14,148,860 instances, these small effect sizes suggest that the practical significance of the differences between groups may be limited, despite the statistical significance indicated by the p-value (Figure 3).

**Figure 3.**
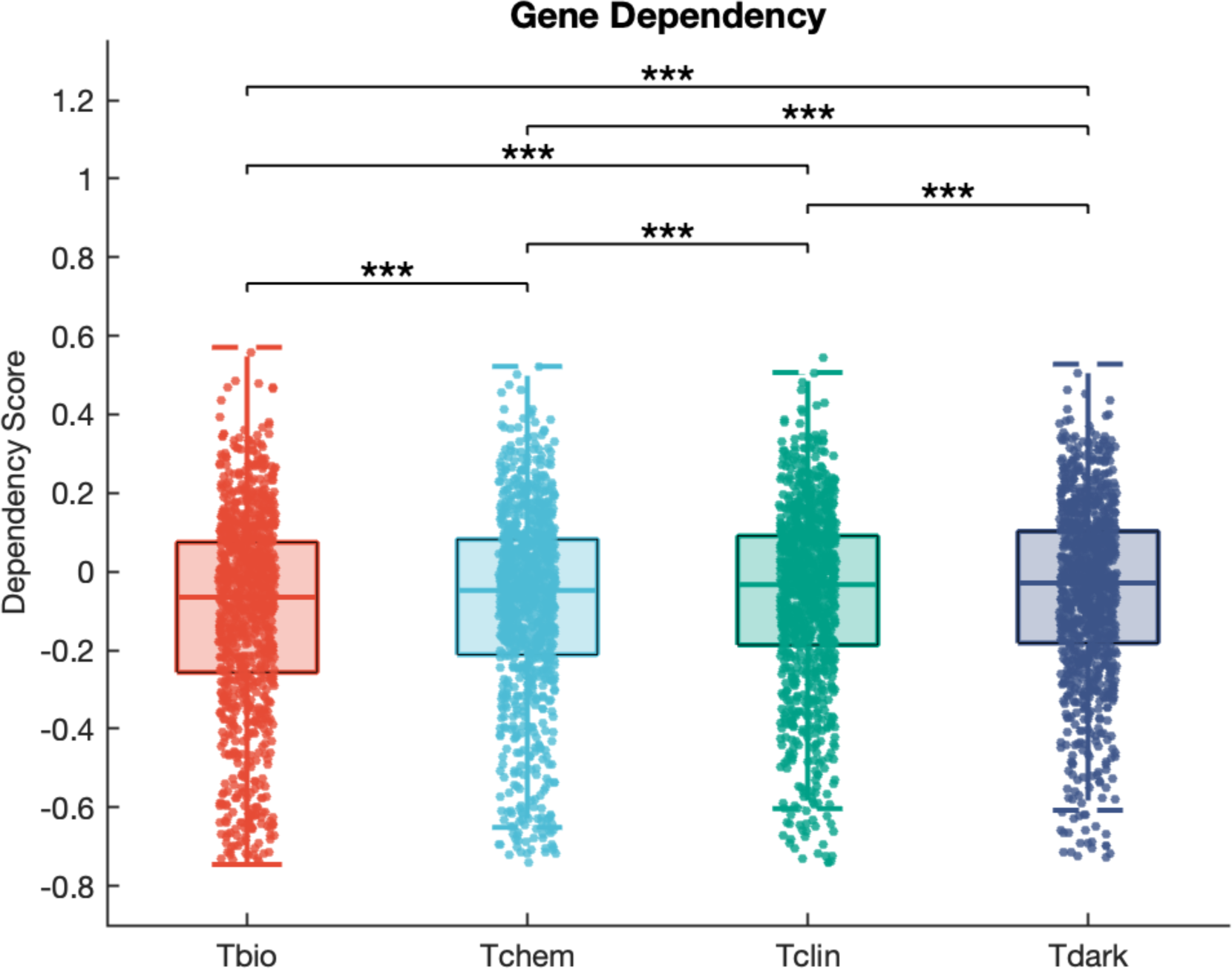
Comparison of gene dependency scores among TDLs in cancer cell lines from the Achilles project. Boxplots show the gene dependency scores corresponding to the Tbio, Tchem, Tclin, and Tdark target development levels. On each box, the central mark indicates the median, and the bottom edge represents the 25th percentile, whereas the top edge of the box represents the 75th percentile. The whiskers extend to the most extreme data points that are not considered outliers. The scatter points within each box plot show the overall distribution of the data points.

### Correlation between gene essentiality and mRNA expression of dark genes

The gene essentiality signature of cell lines has been reported to be related to their mRNA transcription signature^46^. Therefore, we sought to assess the impact of the common essential dark and light genes frequently mutated in cancer on cell fitness. To this end, we examined the correlation between mRNA expression and dependency scores. Among the top genes with the most significant correlation, we found that, among dark genes, the brain cancer cell lines showed significantly greater dependency on the dark gene *NRDE2* than other cell lines (p < 1 x 10^-300^); Figure 4a). Furthermore, cell lines with *NRDE2* mutations exhibited a greater dependency on *NRDE2* expression than other cell lines (p < 1 × 10^-300^). Furthermore, pancreatic cancer cell lines showed significantly greater dependency on the dark gene *WDR7* than other cell lines (p < 1 x 10^-300^); Figure 4b). Additionally, cell lines with *WDR7* mutations exhibited a slightly reduced dependency on *WDR7* expression than other cell lines (p < 1 × 10^-300^). This pattern of dependence was similar to that observed for many light genes, including *CRNKL1* in breast cancer cell lines (Figure 4c) and *MED14* in leukaemia cell lines (Figure 4d). These findings suggest that both dark and light genes play significant roles in cell fitness and may contribute to the development or progression of specific cancer types (Supplementary File 2). Consequently, these results provide valuable insights into the genetic mechanisms involved in these cancers and suggest that these genes could serve as potential targets for the development of novel therapies. Therefore, the CRISPR-based gene editing technology used in the Achilles project has the potential to provide valuable insights for developing new cancer treatments and ultimately improving outcomes for cancer patients^32,47–55^.

**Figure 4.**
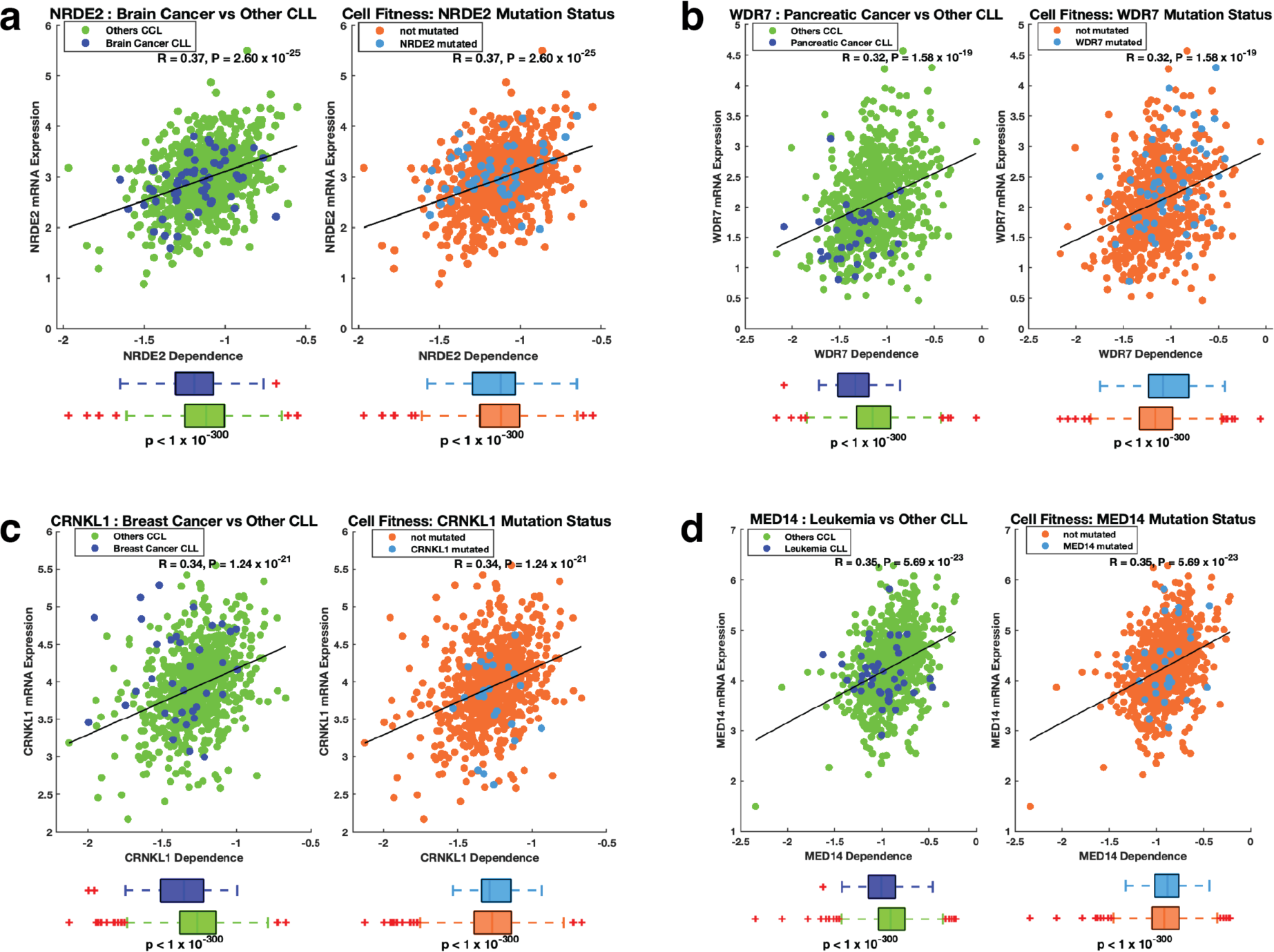
Relationship between gene essentiality and mRNA expression. **a.** From left to right: correlation between *NRDE2* transcript levels and *NRDE2* gene dependence scores, the mean difference in *NRDE2* dependence between breast cancer cell lines and all other cancer cell lines, and the mean difference in *NRDE2* dependence score between *NRDE2* mutant cell lines and cell lines that do not harbour *NRDE2* mutations. **b.** (Left) Correlation between *WDR7* transcript levels and *WDR7* dependence scores, and the mean difference in *WDR7* dependence scores between lymphoma cell lines and all other cancer cell lines. (Right) The mean difference in the *WDR7* dependence score between *WDR7* mutant cell lines and cell lines that did not harbour *WDR7* mutations. **c.** (Left) correlation between *CRNKL1* transcript levels and *CRNKL1* dependence scores, and the mean difference in the *CRNKL1* dependence score between ovarian cancer cell lines and all other cancer cell lines. (Right) The mean difference in the *CRNKL1* dependence score between *CRNKL1* mutant cell lines and cell lines that did not harbour *CRNKL1* mutations. **d** (Left) correlation between *MED14* transcript levels and *MED14* dependence scores, and the mean difference in the *MED14* dependence score between neuroblastoma cell lines and all other cancer cell lines. (Right) The mean difference in the *MED14* dependence score between *MED14* mutant cell lines and cell lines that do not harbour *MED14* mutations.

We further hypothesised that common essential genes are likely to be highly expressed in cancer cell lines. Therefore, we compared the mean transcript levels between the common essential genes and other genes and found that the common essential genes are indeed significantly more highly expressed (Welch t-test; t = 709.7; p < 1 × 10^-300^); see Supplementary Figure 2) in each target development level. Overall, these findings suggest that the “common essential” genes may be a potential target for cancer treatment and further research is warranted to explore this possibility.

### Dark and light genes similarly impact the chemosensitivity of cancer cells

The sensitivity of cancer cell lines to pathway inhibitors is influenced by various factors, such as genetic mutations^56,57^, the targeted pathway^58–61^, the specific inhibitor used^62,63^, and the level of dependence on targeted pathway components^46^. Therefore, we investigated whether CRISPR-derived measures of cellular dependence on pathway components correlate with the response of cell lines to existing drug molecules that inhibit these components, which could inform the identification of optimal drug targets within the pathways (see “Methods” section).

This study classified cancer cell lines from the GDSC database into two categories based on their CRISPR-derived dependency on genes: one group with a higher CRISPR-derived dependency and the other with a lower dependency. We then compared the mean dose responses of 397 pathway inhibitors (Supplementary Figure 3a, also see Supplementary File 3 for the list of inhibitors) between these two groups of cancer cell lines, and only significant results were obtained (p-value < 0.05). We found that 164,225 cases met this criterion, indicating a significant association between pathway inhibitors and the response of cancer cell lines. Notably, these instances encompassed both light genes (Tbio [n = 114,660], and Tchem [n = 21,066]), Tclin [n = 7,173] and dark genes (Tdark [n = 21,326]) (see Supplementary Figure 3c and Supplementary File 3). Among the cell lines dependent on dark genes, we found that the top three inhibitors with the most significant efficacy differences between the groups were cediranib (p = 5.3 × 10^-19^), sepantronium bromide (p = 2.8 × 10^-17^), and KRAS (G12C) inhibitor-12 (p = 1.9 × 10^-16^; Figure 5). These findings suggested that dark genes play a significant role in influencing specific cancer responses to these drugs. Regarding the light genes, the top three inhibitors that demonstrated notable differences in efficacy were nutlin-3a (-) (p = 7.9 × 10^-58^; *MDM2* (Tchem)), rTRAIL (p = 9.8 × 10^-32^; *PLAGL2* (Tbio)) and UNC0638 (p = 2.2 × 10^-28^; *IRF4* (Tbio); see Supplementary Figure 4). Similarly, among the Tclin genes, the top three inhibitors that demonstrated notable differences in efficacy were afatinib (p = 8.9 × 10^-25^ (*ERBB2*) and p = 3.0 × 10^-22^ (*EGFR*)), SNX-2112 (p = 4.6 × 10^-20^) and daporinad (p = 4.7 × 10^-20^ see Supplementary Figure 5). These findings highlight the potential of using dependence scores and drug responses from GDSC to identify additional drug targets in cancer cell lines. Specifically, the Tclin genes associated with drugs whose dose response profiles vary can serve as potential targets for further exploration and therapeutic intervention. Additionally, we found that cell lines with high dependency on pathway genes showed better responses to pathway inhibitors than those with lower dependency (Supplementary Figure 3e).

**Figure 5.**
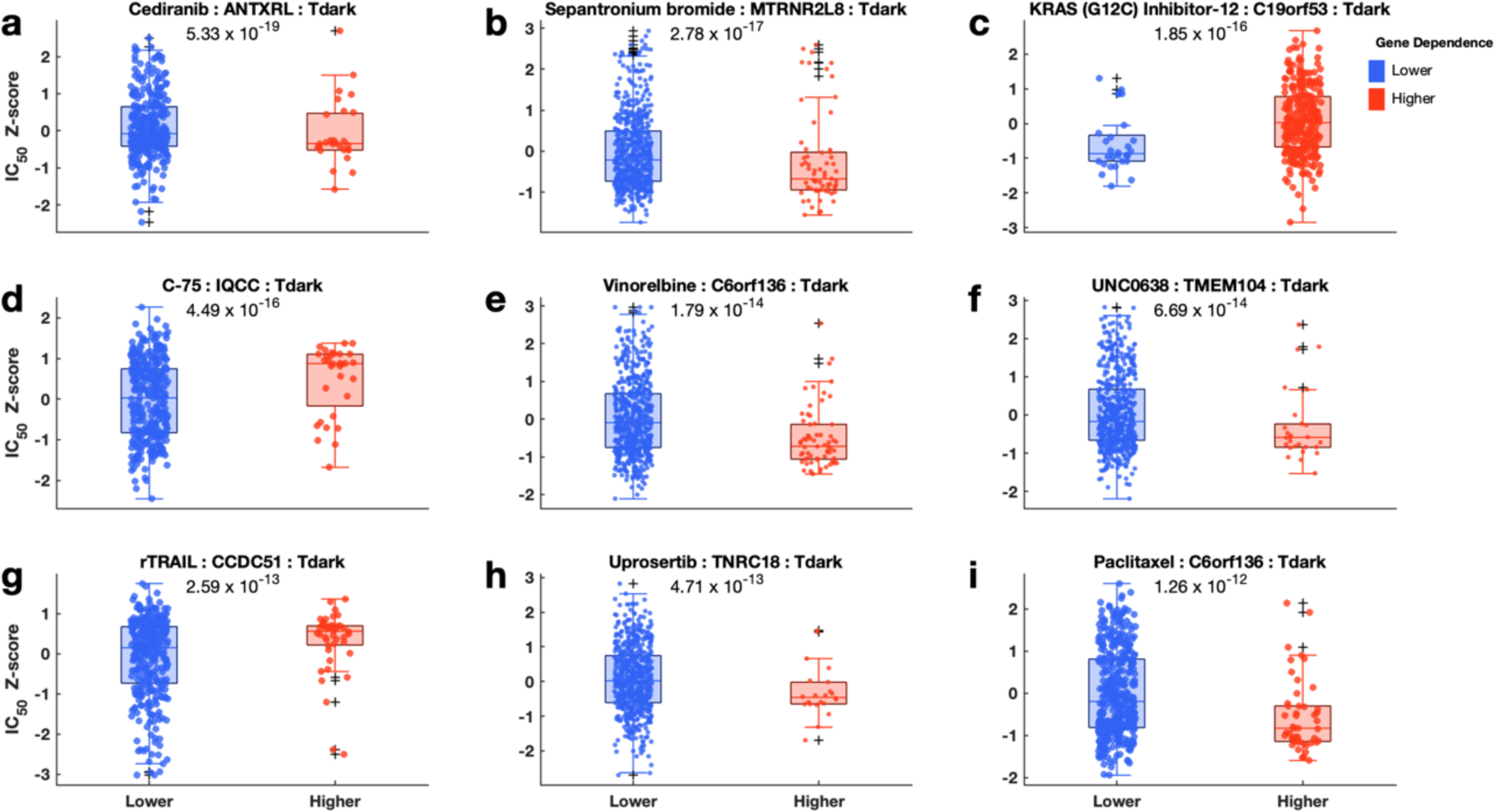
Relationship between Achilles gene dependence scores and the responses of cell lines to pathway inhibitors for Tdark genes. Comparison of the dose-response profiles to pathway inhibitors (**a**: Cediranib, **b**: Sepantronium bromide, **c**: KRAS(G12C) Inhibitor-12, **d**: C-75, **e**: Vinorelbine, **f**: UNC0638, **g**: rTRAIL, **h**: Uprosertib, **i**: Paclitaxel) between the cancer cell lines with lower dependence (boxplots coloured blue) on signalling pathway and those with higher dependence (boxplots coloured red) on signalling pathway. Boxplots show logarithm-transformed mean IC50 values of the cancer cell lines of each group. On each box, the central mark indicates the median, and the bottom edge represents the 25th percentile, whereas the top edge of the box represents the 75th percentile. The whiskers extend to the most extreme data points not considered outliers, and the outliers are plotted individually using the ‘+‘ symbol. The scatter points within each box plot show the overall distribution of the data points.

This study compared the mean dose responses of 397 pathway inhibitors (Supplementary Figure 3b, also see Supplementary File 3 for the list of inhibitors) between cancer cell lines with and without specific gene mutations, and only significant results were obtained (p-value < 0.05). We identified 143,443 instances that met this criterion, indicating a significant association between the pathway inhibitors and the response of the cancer cell lines. Notably, these instances encompassed both light genes (Tbio [n = 92,931], Tchem [n = 18,191]), and Tclin [n = 8,039] and dark genes (Tdark [n = 24,282]) (see Supplementary Figure 3d and Supplementary File 3). Among the cell lines with mutated dark genes, we found that the top three inhibitors with the most significant efficacy differences between groups include vinorelbine (p = 1.5 × 10^-9^), daporinad (p = 2.2 × 10^-9^), and Wee1 inhibitor (p = 4.7 × 10^-9^; Figure 6). These findings highlight the significant role of dark gene mutations in influencing specific cancer responses to these drugs. Additionally, the top three inhibitors associated with light genes included nutlin-3a (-) (p = 2.4 × 10^-38^; *TP53* (Tchem)), paclitaxel (p = 1.8 × 10^-14^; *PLEKHA5* (Tbio)) and uprosertib (p = 6.5 × 10^-12^; *PTEN* (Tchem)) (see Supplementary Figure 6). Furthermore, our study revealed that cancer cells with mutations in specific pathway genes are more responsive to pathway inhibitors than those without mutations (Supplementary Figure 3f).

**Figure 6.**
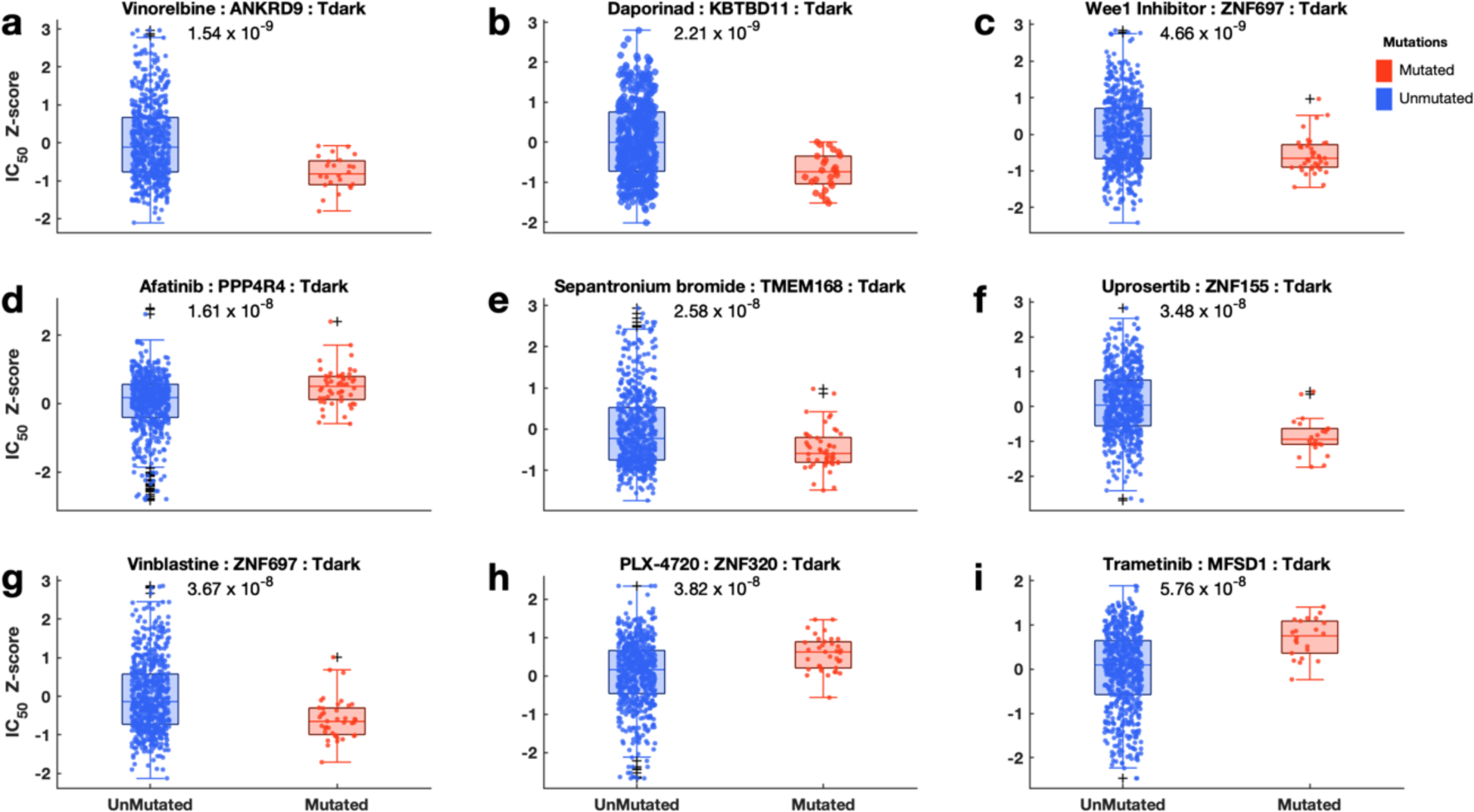
Relationship between mutations of different pathway genes and the responses of the cell line to pathway inhibitors for Tdark genes. Comparison of the dose-response profiles to pathway inhibitors (**a**: Vinorelbine, **b**: Daporinad, **c**: Wee 1 Inhibitor, **d**: Afatinib, **e**: Sepantronium bromide, **f**: Uprosertib, **g**: Vinblastine, **h**: PLX-4720, **i**: Trametinib) between the cancer cell lines without mutation (boxplots blue) on signalling pathway and those with mutation (boxplots coloured red) on signalling pathway. Boxplots show logarithm-transformed mean IC50 values of the cancer cell lines of each group. On each box, the central mark indicates the median, and the bottom edge represents the 25th percentiles, whereas the top edge of the box represents the 75th percentile. The whiskers extend to the most extreme data points not considered outliers, and the outliers are plotted individually using the ‘+‘ symbol. The scatter points within each box plot show the overall distribution of the data points.

Using a chi-squared test of independence, we examined the potential relationship between the target development level of genes and drug sensitivity in cancer cell lines, obtaining a p-value of 0.2133. The results indicated a lack of statistical significance in the observed association, suggesting that mutations in no specific gene class had a biased effect on the drug sensitivity of cancer cells.

We further analysed the response of the cell lines to inhibitors targeting 24 different signalling pathways (Supplementary File 3). Interestingly, we observed that some cell lines with CRISPR-derived dependency scores for specific pathway genes were sensitive to inhibitors of more than one pathway. For example, cell lines dependent on *MEF2C, RUNX1*, and *ALAD* were sensitive to 80%, 44%, and 44% respectively, of the PI3K/mTOR signalling inhibitors. Similarly, these cell lines also displayed sensitivities to 72%, 76%, and 40%, respectively, of the RTK pathway inhibitors profiled by GDSC (also see Supplementary File 3). Furthermore, we found that cell lines dependent on *TUBB4B* and *KLF5* showed resistance to PI3K/mTOR signalling inhibitors (44% and 40% respectively) as well as to RTK pathway inhibitors (64% and 68% respectively), as profiled by the GDSC (see Supplementary File 3). Additionally, we observed that dark gene-dependent cell lines exhibited notable sensitivity to pathway inhibitors, particularly in the presence of mutations. For example, in cell lines with mutations in the PI3K/MTOR signalling pathway, we found that 14 of the top 50 genes were dark genes, including *NPVF*, *FAM122C*, and *TMEM144*, which were associated with mixed responses (i.e., significantly increased sensitivity to some of the inhibitors and significantly decreased sensitivity to others) (Figure 7b). Whereas cell lines with mutations in the RTK pathway, 10 were dark genes, including *ANKRD39, OR7D2,* and *CCDC43*, (7 associated with mixed response, 2 associated with resistance and 1 associated with increased sensitivity) (see Supplementary Figure 7b).

**Figure 7.**
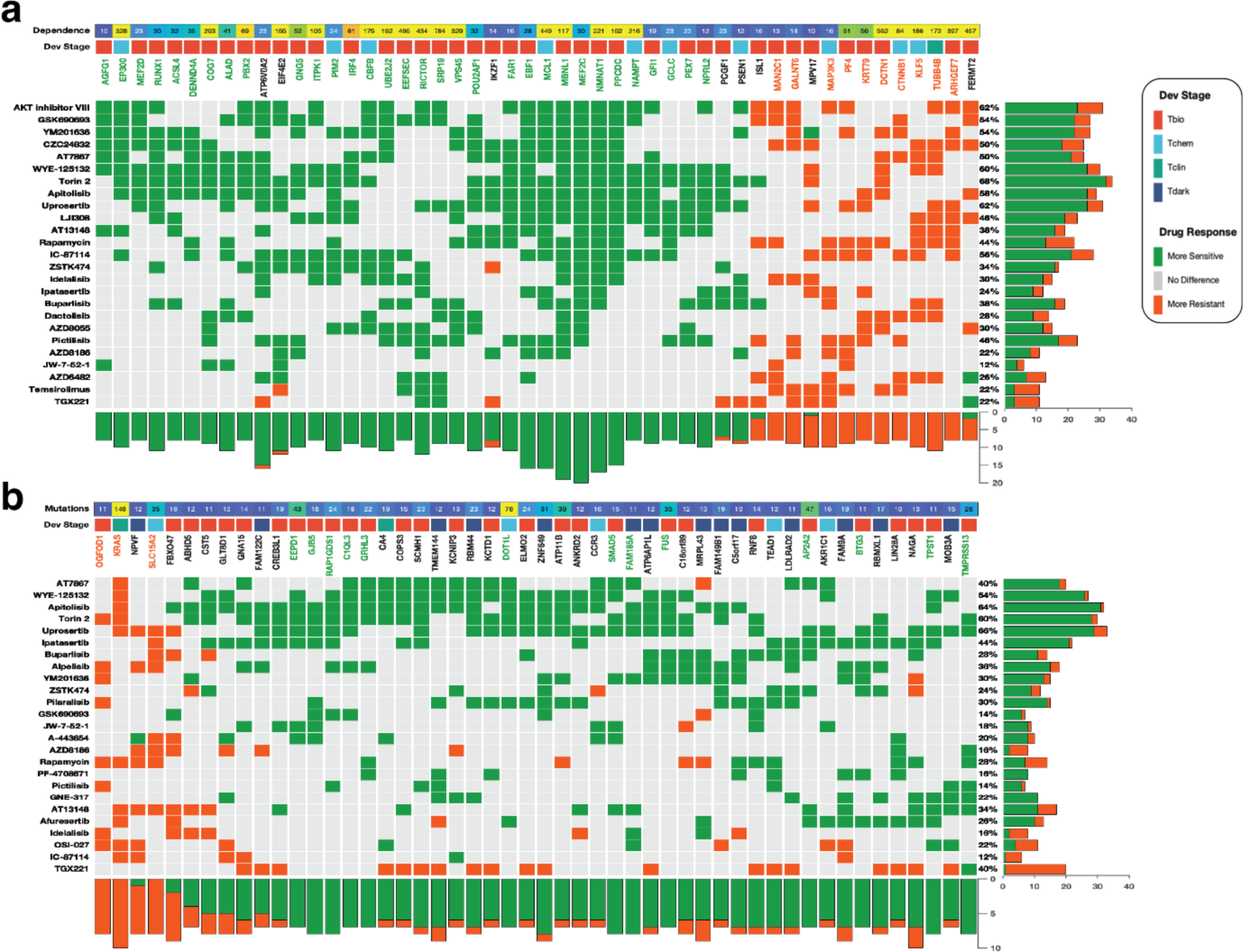
PI3K/MTOR pathway. The relationship between gene dependencies (**a**) or mutations (**b**) and drug responses across cancer cell lines in the PI3K/MTOR pathway. From top to bottom, panels indicate: Dependence; the overall CRISPR-derived gene dependence scores of the gene along that column (**a**). Overall mutation frequencies observed for the gene along the columns (**b**). Clustered heatmap; The marks on the heatmap are coloured based on how a high dependence on, or mutations in, the gene along each column affect the efficacy of the drug given along each row: (1) with green denoting significantly (10% false discovery rate) increased sensitivity, (2) grey for no statistically significant difference between cell line with a higher and lower dependence on the gene, or cell line with mutation in a gene and (3) orange denoting significantly increased resistance (for gene dependence and for gene mutations). The gene names (column labels) are coloured based on the overall calculated effect that high dependence on the gene has on the efficacy of the drug given along rows. Green: all the cell lines are significantly more sensitive to all the pathway inhibitors, orange; all the cell lines are significantly more resistant to all the pathway inhibitors, and black; a mixed response to pathway inhibitors. The bar graphs represent the total number of drugs whose dose-response is significantly increased (green) or decreased (orange).

Our findings highlight that CRISPR-derived estimates of the dependency on signalling pathway components can predict the responsiveness of different primary tumour types to pathway inhibitors. This information could be used to develop targeted therapies for cancers with dark gene mutations or a great dependence on dark genes, which may be more sensitive to pathway inhibitors and respond better to treatment.

### Dark genes and their impact on cancer patients’ survival

We aimed to evaluate the aggressiveness of tumours in patients with genetic alterations in dark genes. We conducted an analysis of disease-free survival (DFS) and overall survival (OS) in cancer patients, considering the mutation status and mRNA expression levels of dark genes to ascertain if specific patient groups exhibited distinct clinical outcomes. We obtained and analysed a pan-cancer dataset from TCGA, which included mRNA expression levels, mutations, and completely de-identified clinical information (refer to the Methods section).

We categorised patients’ tumours into two groups based on their mutation status to assess their impact on OS and DFS. The first group, “tumours with Tdark mutation”, comprised tumours with mutations in dark genes (3,887 samples), while the second group, “tumours without Tdark mutation,” consisted of tumours lacking mutations in dark genes (5,463 samples). We investigated whether the two groups were associated with different clinical outcomes. Using the Kaplan-Meier method^64^, we observed that patients with tumours harbouring dark gene mutations had significantly shorter OS periods (OS = 93.83 months) than those with tumours lacking dark gene mutations (OS = 126.18 months; Figure 8a) (log-rank test; p = 0.002). We also found that DFS periods were significantly shorter (log-rank test; p = 0.008) in patients with tumours containing dark gene mutations than in those without dark gene mutations (Figure 8b).

**Figure 8.**
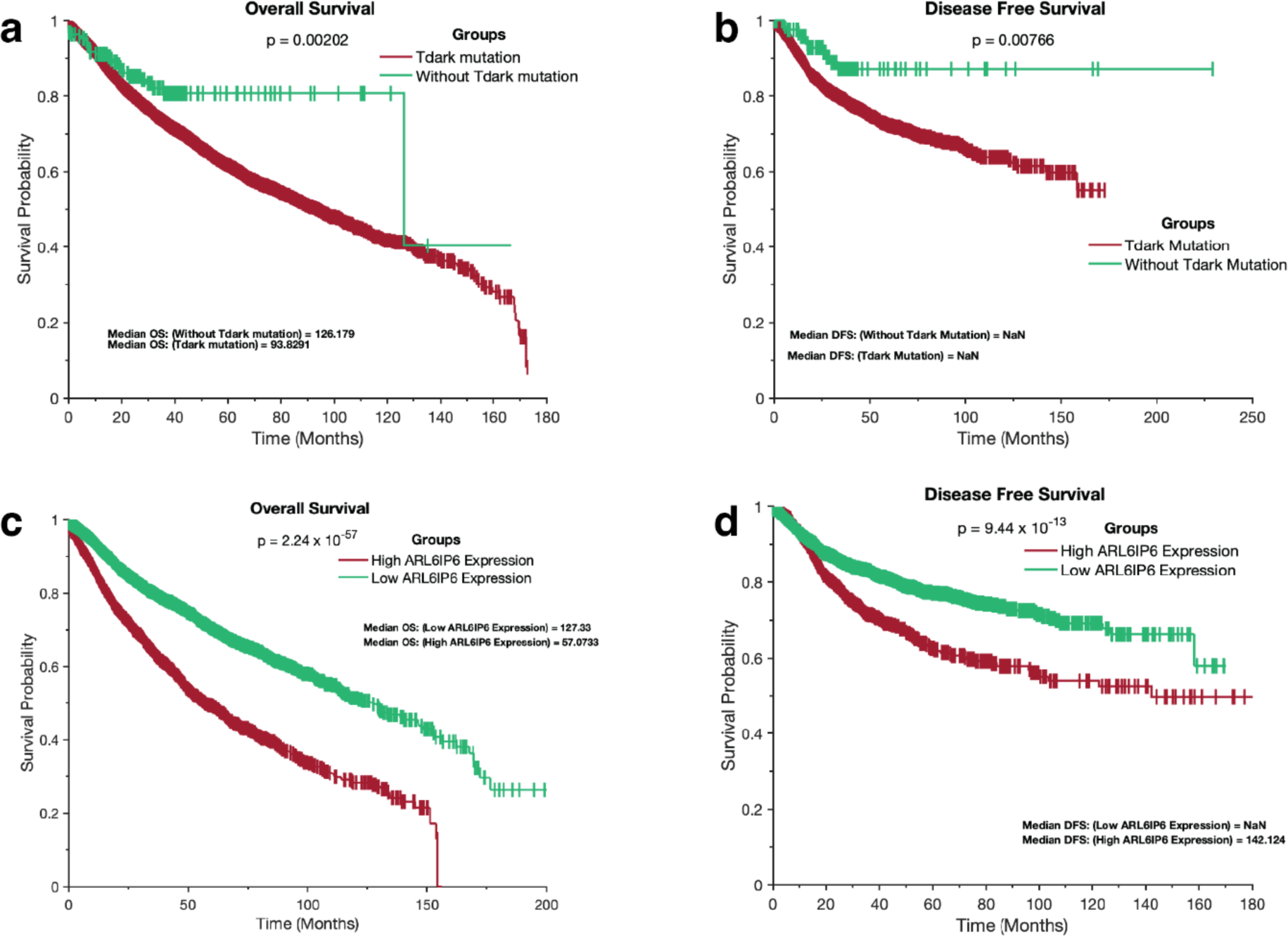
Kaplan–Meier survival curves depicting the impact of dark genes on the survival of patients with cancer. Overall survival periods (**a)** and disease-free survival periods (**b)** of TCGA patients with tumours that had Tdark mutations and tumours without Tdark mutations. Kaplan-Meier curve of the overall survival periods (**c)** and disease-free survival periods (**d)** of TCGA patients with tumours that expressed high and low *ARL6IP6* transcript levels.

To analyse OS and DFS based on mRNA expression levels, we employed the z-score normalisation method to segregate patients’ tumours into two groups: those with higher expression levels of a specific dark gene and those with lower expression levels of a specific dark gene (refer to the Methods section). These groups were defined as high or low expression according to the mRNA transcript levels of a particular dark gene. Using the Kaplan-Meier method, we found that, of the 3,347 dark genes whose expression varied across 8,395 patient tumours analysed, OS periods were significantly shorter for 978 (29.22%) dark genes in patients with tumours expressing high-mRNA transcripts of a specific dark gene. In comparison, OS periods were significantly shorter for 1,258 (37.59%) dark genes in patients with tumours expressing low-mRNA transcripts of a specific dark gene (see Supplementary File 4). DFS analysis revealed that periods were significantly shorter for 755 (22.56%) dark genes in patients with tumours expressing high-mRNA transcripts of a specific dark gene, whereas DFS periods were significantly shorter for 1,011 (30.21%) dark genes in patients with tumours expressing low-mRNA transcripts of a specific dark gene (see Supplementary File 4).

Among the dark genes, we found that patients with tumours with high expression of *ARL6IP6* exhibited the shortest significant OS duration (OS = 57.07 months) compared to those with low gene expression (OS = 127.33 months; Figure 8c). We observed that DFS periods were significantly shorter (log-rank test; p = 9.44 x 10^−13^) for patients with tumours expressing high *ARL6IP6* levels than those with low *ARL6IP6* expression (Figure 8d). Additionally, patients with tumours expressing low *RAI2* levels had the shortest significant OS duration (OS = 68.48 months) compared to those with high expression of this gene (OS = 174.84 months; see Supplementary Figure 8a). We observed that DFS periods were significantly shorter (log-rank test; p = 6.75 x 10^−19^ for patients with tumours expressing low *RAI2* levels than those with high *RAI2* expression (see Supplementary Figure 8b).

Moreover, we investigated the association between the mRNA expression levels of specific dark genes and the aggressiveness of disease in each cancer type and across all the cancer types. We found that, of the top 50 dark genes that exhibited a significant association with reduced OS in each of the 28 cancer types (p-value < 0.05), many genes demonstrated an impact on OS across multiple cancer types. For instance, *ABCA11P* is associated with reduced OS in both oesophageal carcinoma and glioblastoma multiforme. At the same time, *GAGE1* showed a similar effect in five cancer types, including kidney chromophobe, kidney renal papillary cell carcinoma, liver hepatocellular carcinoma, lung squamous cell carcinoma, and prostate adenocarcinoma (see Supplementary Figure 9 and Supplementary File 4). Notably, a significant proportion of genes (694 [17%]) was associated with reduced OS in three different cancer types (see Supplementary Figure 10a). Additionally, our analysis revealed that the cancer types with the highest number of significant genes were LGG (Brain lower grade glioma) and KIRC (Kidney renal clear cell carcinoma), 1,251 and 1,135 genes, respectively (see Supplementary Figure 10b and Supplementary File 4).

In summary, our survival analyses demonstrated that patients with tumours harbouring mutations in the dark genes had significantly shorter overall survival (OS) and disease-free survival (DFS) than those without such mutations. Similarly, a considerable proportion of dark genes showed that patients with tumours expressing high mRNA transcripts had significantly shorter OS and DFS periods than those with low mRNA transcripts. These findings indicate that both the mutation status and mRNA expression levels of dark genes may be useful prognostic markers for cancer patients. Moreover, our study unveiled dark genes whose mRNA expression levels are associated with the aggressiveness of the disease, both in particular cancer types and across multiple cancer types. The identification of these genes provides valuable insights into potential targets for therapeutic interventions and highlights the interconnectedness of specific genes across different cancer types.

## Discussion

In this study, we investigated the potential roles of Tdark genes in cellular processes, cancer development, and progression, and their possible use as drug targets. We observed that Tdark genes constituted a substantial proportion (27.9%) of the genome, highlighting the need for further research to elucidate their function. Our findings also revealed that only a small percentage (3.4%) of Tclin genes are currently utilised as drug targets, indicating a significant potential for developing new drugs targeting relatively well-studied Tchem genes^35,36^.

We identified an increase in the number of publications, antibody counts, and monoclonal antibody counts for essential genes compared with non-essential genes, suggesting a focus on essential genes in research^65–70^. However, we also found that Tdark proteins had the least amount of associated data, indicating a knowledge deficit^27,29,30,36^. Furthermore, our study revealed that Tdark genes are mutated as frequently as light genes in human cancers, with Tdark genes being mutated in 98.96% of all cancers analysed, suggesting significant roles in cancer development and progression.

Our investigation identified *PKHD1L1* as the most frequently mutated Tdark gene across 28 types of cancer, indicating its potential as a broad therapeutic target. Notably, previous research^71–76^ has also indicated the substantial involvement of *PKHD1L1* in cancer, underscoring its probable significance in the development and progression of this disease. We also reported differences in the mean gene dependency scores among Tclin, Tchem, Tbio, and Tdark genes, suggesting that Tdark genes are also essential for the survival of the cancer cell lines and may be promising therapeutic targets^29,32,77^. Furthermore, our study identified critical genes that may be involved in the development and progression of brain cancer, pancreatic cancer, breast cancer, and leukaemia. Therefore, this study sheds light on the genetic mechanisms underlying these cancers, indicating that the identified genes hold promise as potential therapeutic targets.

We found that cancer cell lines with high dependency on pathway genes showed better responses to pathway inhibitors than those with lower dependency. Additionally, cancer cell lines with mutations in specific pathway genes were more responsive to pathway inhibitors than those without mutations, in agreement with a previous study^46^. These findings have important implications for the development of targeted therapies and personalised medical approaches for cancer treatment. Moreover, our findings confirm previous reports that nutlin-3a exhibits sensitivity in cancer cell lines that are dependent on *MDM2*, while demonstrating resistance in cancer cell lines with *TP53* mutations^78,79^. These results underscore the significance of considering the molecular characteristics of cancer cells, including *MDM2* dependence and *TP53* mutation status, when assessing the potential effectiveness of nutlin-3a as a therapeutic intervention.

Furthermore, our study revealed a significant association between mutations in Tdark genes and poorer clinical outcomes in terms of overall survival (OS) and disease-free survival (DFS) in cancer patients. In addition, patients with tumours harbouring Tdark gene mutations had significantly shorter OS and DFS periods than those without such mutations, suggesting that mutations in Tdark genes may be important predictors of clinical outcomes. Moreover, our analysis revealed a strong correlation between the expression of Tdark genes and OS and DFS. Notably, we observed that high expression of *ARL6IP6* and low expression of *RAI2* were particularly associated with the shortest OS and DFS periods. It is worth noting that previous studies have shown that decreased *RAI2* expression is linked to poor prognosis in colorectal cancer^80,81^ and breast cancer^82–84^. These findings underscore the potential significance of Tdark as a valuable prognostic marker in cancer.

In conclusion, our findings underscore the importance of incorporating genetic information into cancer treatment and highlight the potential of personalised medicine approaches. Furthermore, the results demonstrated that Tdark genes are important players in cancer development, warranting further research into their biological functions and potential as targets for cancer therapy.

## Methods

We analysed a dataset comprising 10,528 patient-derived tumours representing 32 distinct human cancers, obtained from cBioPortal^31^ version 3.1.9 (http://www.cbioportal.org). The acquired data included somatic gene mutations (point mutations and small insertions/deletions), mRNA expression, and comprehensive de-identified clinical data. We further filtered the dataset to include only cancer studies with clinical information for profiled patients. The final datasets encompassed 8,395 patient-derived tumour samples representing 28 distinct human cancer types (see Supplementary File 1 for cancer study details).

### Distribution and research focus on light and dark genes across the target development levels (TDLs)

We obtained human gene classifications based on target development levels (TDLs) from the Pharos^29^ interface (version 3.15.1) (https://pharos.nih.gov/). The IDG project classified proteins into four TDLs based on the level of clinical, biological, and chemical investigations conducted on each protein. These TDLs include light genes (Tbio, Tclin, and Tchem) and dark genes (Tdark)^85^:

- Tclin proteins are drug targets associated with at least one approved drug, and their mechanism of action (MoA) is known^29,35^.
- Tchem proteins do not have established connections to approved drugs based on MoA. However, they are recognised for their exceptional ability to bind with high potency to small molecules, surpassing the bioactivity cutoff values: ≤30 nM for kinases, ≤100 nM for GPCRs and nuclear receptors, ≤10 μM for ion channels, and ≤1 μM for other target families^29,85^.
- Tbio are proteins with well-studied biology and meet certain criteria, such as having a fractional publication count above 5, being annotated with a Gene Ontology Molecular Function or Biological Process with an Experimental Evidence code and having confirmed OMIM phenotype(s)^29,35^.
- Tdark are understudied proteins that do not meet the criteria for the above three categories^35^. They meet at least two of the following conditions: A PubMed text-mining score <5, <=3 Gene RIFs or <=50 Antibodies available per Antibodypedia^85^.

We analysed a dataset of 20,412 targets: light genes (Tbio [n = 12,058], Tclin [n = 704], Tchem [n = 1,971]) and dark genes (Tdark [n = 5,679]). Additionally, we compared gene distribution, and TDLs between TCRD versions 4.3.4 and 6.13.4 to investigate gene classification changes over time. We also gathered information on antibody counts, monoclonal antibody counts, and PubMed publication counts from Pharos for further insight into the research focus within each TDL.

### Determination of the extent to which dark genes are mutated in human cancer types

To assess the extent of dark gene mutations in cancer compared to light genes, we integrated TDL-annotated protein-coding gene information from the Pharos database with the TCGA project^31^ dataset of 8,395 patient-derived tumours representing 28 distinct human cancers. We determined the overall mutation frequencies in genes of each developmental level (Tdark, Tclin, Tchem, and Tbio) across the tumours of each cancer type and all cancer types. These analyses enabled us to understand how dark genes are altered within and across the 28 most common human cancer types.

### Assessment of the impact of Tdark genes on cancer cell lines

To further evaluate the potential role of dark genes in cancer, we mined datasets from the Achilles project at the DepMap Portal version 21Q1 on the fitness of over 80 cell lines derived from 35 different human cancer types following CRISPR knockouts of 18,333 individual genes. See https://depmap.org/portal/ for information on Achilles’ CRISPR-derived gene dependency descriptions. Briefly, a lower score indicates a higher likelihood of cell line dependency on a given gene. A score of 0 corresponds to a non-essential gene, while −1 corresponds to the median of all “common essential genes”. The database groups genes into four primary categories based on cell line fitness after CRISPR-mediated gene knockouts:

- Common essential genes – consistently ranked in the top X most depleted genes in at least 90% of cell lines.
- Strongly selective genes – genes with a dependency skewed-likelihood ratio test (LRT) value > 100.
- Essential genes – associated with cell fitness in only one or a few cell lines, but with a dependency skewed-LRT value < 100.
- Non-essential genes – show no effect on the cell fitness in any of the 688 tested cell lines.

We incorporated TDL information from Pharos and CRISPR data to examine the impact of Tdark genes on cancer cell lines. Our assessment involved comparing the mean gene dependency scores obtained from CRISPR data to determine the importance of genes within each developmental stage for cancer cell lines across all cancer types and within each cancer type.

Additionally, we calculated and compared the number of common essential and non-essential genes across the Pharos development levels (Supplementary File 1) to understand how gene dependence relates to TDL classes. We then compared the number of PubMed publications, monoclonal antibodies, and polyclonal antibodies available for each group of genes (common essential versus non-essential genes).

### Impact of common essential genes on cell fitness

To further investigate gene essentiality, we collected mRNA transcription data for 756 cancer cell lines from the Cancer Cell Line Encyclopaedia (CCLE) database^34^. Using CRISPR-derived dependence scores, we identified essential genes across cell lines by counting the number of instances in which each gene’s score was less than −0.5. This threshold was chosen based on the recommendation of the Achilles project, indicating reduced cell fitness after CRISPR-mediated gene knockouts. We then tallied instances where the CRISPR-derived dependence score was less than −0.5 for each gene in each cell line to identify genes associated with a reduction in cell fitness across all cell lines. Genes with a significant number of instances were identified as essential, as they were critical for cell survival and growth. Finally, we compared Achilles CRISPR-based fitness scores with transcription profiles.

### Evaluation of the extent to which the dark genome affects the chemosensitivity of cancer cells in relation to the light genome

We analysed drug-response data from the Genomics of Drug Sensitivity in Cancer (GDSC) database^33^ (www.cancerRxgene.org), to investigate the influence of the dark genome on cancer cell chemosensitivity compared to the light genome. The GDSC database provides comprehensive information on human cancer cell lines treated with a wide range of anticancer drugs that target various signalling pathways. We retrieved the dose-responses for 380 cancer cell lines to 397 drugs that target components of 24 different pathways. Additionally, we used Pharos to obtain information on the TDL annotation of light and dark genes.

We conducted a series of analyses for each pathway gene (e.g., *MFSD1*) in the context of the CRISPR dependency scores and mutations in cancer cell lines. Firstly, we divided the cell lines into two groups: (1) those with a high CRISPR dependency score on the gene (e.g., *MFSD1* dependence score < −0.5), and (2) those with low dependence on that gene (e.g., *MFSD1* dependence score > 0.5). Subsequently, we used the Wilcoxon rank-sum test to compare the IC50 values of 397 pathway inhibitors between these two groups of cell lines. Additionally, for each pathway gene, we further categorised the cell lines into two groups: (1) those with mutations in the specific gene (e.g., cell lines with *MFSD1* mutations), and (2) those without mutations in that gene (e.g., cell lines with no *MFSD1* mutations). Next, we compared the logarithm-transformed IC50 values of each anticancer drug between these two groups of cell lines using the Wilcoxon rank-sum test.

By conducting these analyses, we aimed to determine the extent to which dark and light genes affect cancer chemosensitivity and to identify the types of drugs and targeted pathways that are more relevant for tumours with alterations in specific dark and light genes.

### Assessing the impact of dark genome alterations on cancer aggressiveness

To evaluate the influence of dark genome alterations on cancer aggressiveness, we integrated genetic alteration data, including mRNA transcription changes and gene mutations of 28 human cancer types, with clinical outcomes of patients in the TCGA. First, for each cancer type, we segregated patients’ tumours into two groups: 1) those without mutations and 2) mutations across all dark genes, or 1) those expressing higher mRNA transcripts of a particular dark gene and 2) those expressing lower mRNA transcripts of a particular dark gene. We then applied the Kaplan-Meier method and Log-rank test to compare the duration of overall survival (OS) and disease-free survival (DFS) between the two groups of tumours across all cancer types.

By comparing the OS and DFS durations, we sought to determine whether alterations in dark genes influence cancer aggressiveness and, if so, the extent of their impact. Furthermore, we investigated the association between mRNA expression levels of specific dark genes and the aggressiveness of disease in each cancer types and across all the cancer types. This assessment allowed us to identify potential dark gene targets that could have clinical significance in the prognosis and treatment of cancer.

### Statistical analyses

All statistical analyses were performed using MATLAB R2021a software. Where appropriate, we used the independent sample Student t-test, Welch test, the Wilcoxon rank-sum test and the one-way Analysis of Variance to compare groups of continuous variables. Statistical tests were considered significant at p < 0.05 for single comparisons, whereas the p-values of multiple comparisons were adjusted using the Bonferroni correction method.

## Supporting information

Description of Supplementary Files

## Data Availability

The data that support our results are available in this manuscript, the supplementary data, and from the following repositories: Pharos; https://pharos.nih.gov/, The cancer Genome Atlas (TCGA); https://www.cancer.gov/ccg/research/genome-sequencing/tcga, the Genomics of Drug Sensitivity in Cancer; https://www.cancerrxgene.org/, the Cancer Cell Line Encyclopaedia; https://portals.broadinstitute.org/ccle/data and the Project Achilles; https://depmap.org/portal/.

## Ethics approval

The study protocol was approved by The University of Zambia; Health Sciences Research Ethics Committee IRB00011000. The publicly available datasets were collected by the TCGA, CCLE, Achilles, IDG and GDSC projects and made available via their respective project databases. The methods used here were performed following the relevant policies, regulations and guidelines provided by the TCGA, CCLE, DepMap, IDG and GDSC projects.

## Author contributions

### Conceptualisation

Doris Kafita, Kevin Dzobo and Musalula Sinkala.

### Methodology

Doris Kafita, Panji Nkhoma, Kevin Dzobo and Musalula Sinkala.

### Formal analysis

Doris Kafita, Panji Nkhoma, Kevin Dzobo and Musalula Sinkala.

### Visualisation

Doris Kafita, Kevin Dzobo and Musalula Sinkala.

### Writing – original draft

Doris Kafita and Musalula Sinkala.

### Writing – review & editing

Doris Kafita, Panji Nkhoma, Kevin Dzobo and Musalula Sinkala.

## References

1. Chi, K. The dark side of the human Genome. Nature 538, 275–277 (2016).

2. Ezkurdia, I. et al. Multiple evidence strands suggest that theremay be as few as 19 000 human protein-coding genes. Hum Mol Genet 23, 5866–5878 (2014).

3. Pertea, M. et al. CHESS: A new human gene catalog curated from thousands of large-scale RNA sequencing experiments reveals extensive transcriptional noise. Genome Biol 19, (2018).

4. Abascal, F. et al. Loose ends: Almost one in five human genes still have unresolved coding status. Nucleic Acids Res 46, 7070–7084 (2018).

5. Nerenz, R. D. & Lefferts, J. Our genome’s ‘Dark Matter’ is the next frontier in molecular diagnostics. Clinical Chemistry vol. 63 792–793 Preprint at https://doi.org/10.1373/clinchem.2016.268607 (2017).

6. Richards, A. L. et al. Environmental perturbations lead to extensive directional shifts in RNA processing. PLoS Genet 13, (2017).

7. Li, Y., et al. Pathway perturbations in signaling networks: Linking genotype to phenotype. Seminars in Cell and Developmental Biology vol. 99 3–11 Preprint at https://doi.org/10.1016/j.semcdb.2018.05.001 (2020).

8. Kafita, D. et al. High ELF4 expression in human cancers is associated with worse disease outcomes and increased resistance to anticancer drugs. PLoS One 16, (2021).

9. Jackson, M., Marks, L., May, G. H. W. & Wilson, J. B. The genetic basis of disease. Essays in Biochemistry vol. 62 643–723 Preprint at https://doi.org/10.1042/EBC20170053 (2018).

10. Albanaz, A., Rodrigues, C., Pires, D. & Ascher, D. Combating mutations in genetic disease and drug resistance: understanding molecular mechanisms to guide drug design - CORE Reader. Expert Opin Drug Discov 12, 553–563 (2017).

11. Amin, A. R. M. R. et al. Evasion of anti-growth signaling: A key step in tumorigenesis and potential target for treatment and prophylaxis by natural compounds. Seminars in Cancer Biology vol. 35 S55–S77 Preprint at https://doi.org/10.1016/j.semcancer.2015.02.005 (2015).

12. Diederichs, S. et al. The dark matter of the cancer genome: aberrations in regulatory elements, untranslated regions, splice sites, non-coding RNA and synonymous mutations. EMBO Mol Med 8, 442–457 (2016).

13. Zhao, M., Kim, P., Mitra, R., Zhao, J. & Zhao, Z. TSGene 2.0: An updated literature-based knowledgebase for Tumor Suppressor Genes. Nucleic Acids Res 44, D1023–D1031 (2016).

14. Wang, L. H., Wu, C. F., Rajasekaran, N. & Shin, Y. K. Loss of tumor suppressor gene function in human cancer: An overview. Cellular Physiology and Biochemistry vol. 51 2647–2693 Preprint at https://doi.org/10.1159/000495956 (2019).

15. Kontomanolis, E. N. et al. Role of oncogenes and tumor-suppressor genes in carcinogenesis: A review. Anticancer Research vol. 40 6009–6015 Preprint at https://doi.org/10.21873/anticanres.14622 (2020).

16. Chandrashekar, P. et al. Somatic selection distinguishes oncogenes and tumor suppressor genes. Bioinformatics 36, 1712–1717 (2020).

17. Leiserson, M. D. M. et al. Pan-cancer network analysis identifies combinations of rare somatic mutations across pathways and protein complexes. Nat Genet 47, 106–114 (2015).

18. Chaitankar, V. et al. Next generation sequencing technology and genomewide data analysis: Perspectives for retinal research. Progress in Retinal and Eye Research vol. 55 1–31 Preprint at https://doi.org/10.1016/j.preteyeres.2016.06.001 (2016).

19. Kandoth, C. et al. Mutational landscape and significance across 12 major cancer types. Nature 502, 333–339 (2013).

20. Kingsmore, S. F. et al. Measurement of genetic diseases as a cause of mortality in infants receiving whole genome sequencing. NPJ Genom Med 5, (2020).

21. Petrikin, J. E., Willig, L. K., Smith, L. D. & Kingsmore, S. F. Rapid whole genome sequencing and precision neonatology. Seminars in Perinatology vol. 39 623–631 Preprint at https://doi.org/10.1053/j.semperi.2015.09.009 (2015).

22. Pervez, M. T. et al. A Comprehensive Review of Performance of Next-Generation Sequencing Platforms. BioMed Research International vol. 2022 Preprint at https://doi.org/10.1155/2022/3457806 (2022).

23. Nagai, H. & Kim, Y. H. Cancer prevention from the perspective of global cancer burden patterns. Journal of Thoracic Disease vol. 9 448–451 Preprint at https://doi.org/10.21037/jtd.2017.02.75 (2017).

24. Tran, K. B. et al. The global burden of cancer attributable to risk factors, 2010–19: a systematic analysis for the Global Burden of Disease Study 2019. The Lancet 400, 563–591 (2022).

25. Lin, L. et al. Global, regional, and national cancer incidence and death for 29 cancer groups in 2019 and trends analysis of the global cancer burden, 1990–2019. J Hematol Oncol 14, (2021).

26. Stefanoudakis, D. et al. The Potential Revolution of Cancer Treatment with CRISPR Technology. Cancers (Basel) 15, 1813 (2023).

27. Oprea, T. I. Exploring the dark genome: implications for precision medicine. Mammalian Genome vol. 30 192–200 Preprint at https://doi.org/10.1007/s00335-019-09809-0 (2019).

28. Pandey, A. K., Lu, L., Wang, X., Homayouni, R. & Williams, R. W. Functionally enigmatic genes: A case study of the brain ignorome. PLoS One 9, (2014).

29. Oprea, T. I. et al. Unexplored therapeutic opportunities in the human genome. Nature Reviews Drug Discovery vol. 17 317–332 Preprint at https://doi.org/10.1038/nrd.2018.14 (2018).

30. Brown, S. D. M. & Lad, H. V. The dark genome and pleiotropy: challenges for precision medicine. Mammalian Genome 30, 212–216 (2019).

31. Weinstein, J. N. et al. The cancer genome atlas pan-cancer analysis project. Nat Genet 45, 1113–1120 (2013).

32. Hart, T. et al. High-Resolution CRISPR Screens Reveal Fitness Genes and Genotype-Specific Cancer Liabilities. Cell 163, 1515–1526 (2015).

33. Yang, W. et al. Genomics of Drug Sensitivity in Cancer (GDSC): A resource for therapeutic biomarker discovery in cancer cells. Nucleic Acids Res 41, (2013).

34. Ghandi, M. et al. Next-generation characterization of the Cancer Cell Line Encyclopedia. Nature 569, 503–508 (2019).

35. Sheils, T. et al. How to Illuminate the Druggable Genome Using Pharos. Curr Protoc Bioinformatics 69, (2020).

36. Nguyen, D. T. et al. Pharos: Collating protein information to shed light on the druggable genome. Nucleic Acids Res 45, D995–D1002 (2017).

37. Brachova, P. et al. TP53 oncomorphic mutations predict resistance to platinum- and taxane-based standard chemotherapy in patients diagnosed with advanced serous ovarian carcinoma. Int J Oncol 46, 607–618 (2015).

38. Zhang, Y., Cao, L., Nguyen, D. & Lu, H. TP53 mutations in epithelial ovarian cancer. Translational Cancer Research vol. 5 650–663 Preprint at https://doi.org/10.21037/tcr.2016.08.40 (2016).

39. Iwanicki, M. P., et al. Mutant p53 regulates ovarian cancer transformed phenotypes through autocrine matrix deposition. JCI Insight 1, (2019).

40. Oien, D. B. & Chien, J. TP53 mutations as a biomarker for high-grade serous ovarian cancer: Are we there yet? Translational Cancer Research vol. 5 S264–S268 Preprint at https://doi.org/10.21037/tcr.2016.07.45 (2016).

41. Parkinson, C. A. et al. Exploratory Analysis of TP53 Mutations in Circulating Tumour DNA as Biomarkers of Treatment Response for Patients with Relapsed High-Grade Serous Ovarian Carcinoma: A Retrospective Study. PLoS Med 13, (2016).

42. Silwal-Pandit, L., Langerød, A. & Børresen-Dale, A. L. TP53 mutations in breast and ovarian cancer. Cold Spring Harb Perspect Med 7, (2017).

43. Tuna, M. et al. Clinical relevance of TP53 hotspot mutations in high-grade serous ovarian cancers. Br J Cancer 122, 405–412 (2020).

44. Yamulla, R. J., Nalubola, S., Flesken-Nikitin, A., Nikitin, A. Y. & Schimenti, J. C. Most Commonly Mutated Genes in High-Grade Serous Ovarian Carcinoma Are Nonessential for Ovarian Surface Epithelial Stem Cell Transformation. Cell Rep 32, (2020).

45. Wallis, B., Bowman, K. R., Lu, P. & Lim, C. S. The Challenges and Prospects of p53-Based Therapies in Ovarian Cancer. Biomolecules vol. 13 Preprint at https://doi.org/10.3390/biom13010159 (2023).

46. Sinkala, M., Nkhoma, P., Mulder, N. & Martin, D. P. Integrated molecular characterisation of the MAPK pathways in human cancers reveals pharmacologically vulnerable mutations and gene dependencies. Commun Biol 4, (2021).

47. Padayachee, J. & Singh, M. Therapeutic applications of CRISPR/Cas9 in breast cancer and delivery potential of gold nanomaterials. Nanobiomedicine vol. 7 Preprint at https://doi.org/10.1177/1849543520983196 (2020).

48. Zhang, H. et al. Application of the CRISPR/Cas9-based gene editing technique in basic research, diagnosis, and therapy of cancer. Molecular Cancer vol. 20 Preprint at https://doi.org/10.1186/s12943-021-01431-6 (2021).

49. Chira, S. et al. Genome Editing Approaches with CRISPR/Cas9 for Cancer Treatment: Critical Appraisal of Preclinical and Clinical Utility, Challenges, and Future Research. Cells vol. 11 Preprint at https://doi.org/10.3390/cells11182781 (2022).

50. Tufail, M. Genome editing: An essential technology for cancer treatment. Medicine in Omics 4, 100015 (2022).

51. Katti, A., Diaz, B. J., Caragine, C. M., Sanjana, N. E. & Dow, L. E. CRISPR in cancer biology and therapy. Nature Reviews Cancer vol. 22 259–279 Preprint at https://doi.org/10.1038/s41568-022-00441-w (2022).

52. Liu, Z., et al. Recent advances and applications of CRISPR-Cas9 in cancer immunotherapy. Molecular Cancer vol. 22 Preprint at https://doi.org/10.1186/s12943-023-01738-6 (2023).

53. Rabaan, A. A. et al. Application of CRISPR/Cas9 Technology in Cancer Treatment: A Future Direction. Current Oncology vol. 30 1954–1976 Preprint at https://doi.org/10.3390/curroncol30020152 (2023).

54. Meng, H. et al. Application of CRISPR-Cas9 gene editing technology in basic research, diagnosis and treatment of colon cancer. Frontiers in Endocrinology vol. 14 Preprint at https://doi.org/10.3389/fendo.2023.1148412 (2023).

55. McLean, B. et al. A CRISPR Path to Finding Vulnerabilities and Solving Drug Resistance: Targeting the Diverse Cancer Landscape and Its Ecosystem. Advanced Genetics 3, 2200014 (2022).

56. Zhang, N. et al. Predicting Anticancer Drug Responses Using a Dual-Layer Integrated Cell Line-Drug Network Model. PLoS Comput Biol 11, (2015).

57. Wang, Y., Fang, J. & Chen, S. Inferences of drug responses in cancer cells from cancer genomic features and compound chemical and therapeutic properties. Sci Rep 6, (2016).

58. Wang, X., Sun, Z., Zimmermann, M. T., Bugrim, A. & Kocher, J. P. Predict drug sensitivity of cancer cells with pathway activity inference. BMC Med Genomics 12, (2019).

59. Ben-Hamo, R. et al. Predicting and affecting response to cancer therapy based on pathway-level biomarkers. Nat Commun 11, (2020).

60. Zhao, W. et al. Large-Scale Characterization of Drug Responses of Clinically Relevant Proteins in Cancer Cell Lines. Cancer Cell 38, 829–843.e4 (2020).

61. Tang, Y. C. & Gottlieb, A. Explainable drug sensitivity prediction through cancer pathway enrichment. Sci Rep 11, (2021).

62. Parca, L. et al. Modeling cancer drug response through drug-specific informative genes. Sci Rep 9, (2019).

63. Xia, F. et al. A cross-study analysis of drug response prediction in cancer cell lines. Brief Bioinform 23, (2022).

64. Goel, M., Kishore, J. & Khanna, P. Understanding survival analysis: Kaplan-Meier estimate. Int J Ayurveda Res 1, 274 (2010).

65. Zhang, W., Quevedo, J. & Fries, G. R. Essential genes from genome-wide screenings as a resource for neuropsychiatric disorders gene discovery. Transl Psychiatry 11, (2021).

66. Blomen, V. A. et al. Gene essentiality and synthetic lethality in haploid human cells. Science (1979) 350, 1092–1096 (2015).

67. Chen, H. et al. New insights on human essential genes based on integrated analysis and the construction of the HEGIAP web-based platform. Brief Bioinform 21, 1397–1410 (2019).

68. Schonfeld, E., Vendrow, E., Vendrow, J. & Schonfeld, E. On the relation of gene essentiality to intron structure: A computational and deep learning approach. Life Sci Alliance 4, (2021).

69. Caldu-Primo, J. L., Verduzco-Martínez, J. A., Alvarez-Buylla, E. R. & Davila-Velderrain, J. In vivo and in vitro human gene essentiality estimations capture contrasting functional constraints. NAR Genom Bioinform 3, (2021).

70. Cacheiro, P. & Smedley, D. Essential genes: a cross-species perspective. Mammalian Genome Preprint at https://doi.org/10.1007/s00335-023-09984-1 (2023).

71. Kohrs, B. Exploring the Role of the PKHD1L1 Gene in Epithelial Cancer Cells. Denison Student Scholarship. 47 (2021).

72. Zheng, C., Quan, R., Xia, E. J., Bhandari, A. & Zhang, X. Original tumour suppressor gene polycystic kidney and hepatic disease 1-like 1 is associated with thyroid cancer cell progression. Oncol Lett 18, 3227–3235 (2019).

73. Wang, Y. et al. Screening and identification of biomarkers associated with the diagnosis and prognosis of lung adenocarcinoma. J Clin Lab Anal 34, (2020).

74. Han, L. K. et al. Identification of prognostic genes in lung adenocarcinoma immune microenvironment. Chinese Medical Journal vol. 134 2125–2127 Preprint at https://doi.org/10.1097/CM9.0000000000001367 (2021).

75. Al-Dherasi, A. et al. A seven-gene prognostic signature predicts overall survival of patients with lung adenocarcinoma (LUAD). Cancer Cell Int 21, (2021).

76. Yang, Y. et al. Excavation of diagnostic biomarkers and construction of prognostic model for clear cell renal cell carcinoma based on urine proteomics. Front Oncol 13, (2023).

77. Dickinson, M. E. et al. High-throughput discovery of novel developmental phenotypes. Nature 537, 508–514 (2016).

78. Crane, E. K. et al. Nutlin-3a: A potential therapeutic opportunity for TP53 wild-type ovarian carcinomas. PLoS One 10, (2015).

79. Walter, R. F. H. et al. Inhibition of MDM2 via Nutlin-3A: A Potential Therapeutic Approach for Pleural Mesotheliomas with MDM2-Induced Inactivation of Wild-Type P53. J Oncol 2018, (2018).

80. Yan, W. et al. Retinoic acid-induced 2 (RAI2) is a novel tumor suppressor, and promoter region methylation of RAI2 is a poor prognostic marker in colorectal cancer. Clin Epigenetics 10, (2018).

81. Zhang, W. et al. Retinoic Acid-Induced 2 (RAI2) Is a Novel Antagonist of Wnt/β-Catenin Signaling Pathway and Potential Biomarker of Chemosensitivity in Colorectal Cancer. Front Oncol 12, (2022).

82. Esposito, M. & Kang, Y. RAI2: Linking retinoic acid signaling with metastasis suppression. Cancer Discov 5, 466–468 (2015).

83. Werner, S. et al. Suppression of early hematogenous dissemination of human breast cancer cells to bone marrow by retinoic acid–induced 2. Cancer Discov 5, 506–519 (2015).

84. Jiao, Y., Li, S., Gong, J., Zheng, K. & Xie, Y. Comprehensive analysis of the expression and prognosis for RAI2: A promising biomarker in breast cancer. Front Oncol 13, (2023).

85. Lin, Y. et al. Drug target ontology to classify and integrate drug discovery data. J Biomed Semantics 8, (2017).

