## Supplementary material for "Shedding Light on the Dark Genome: Insights into the Genetic, CRISPR-based, and Pharmacological Dependencies of Human Cancers and Disease Aggressiveness": Description of Supplementary Files

### Description of Additional Supplementary Files

**Supplementary File 1:** Supplementary data of light and dark genes and cancer studies. The spreadsheet contains the following results/datasets according to the sheet name. **Pharos Data:** The distribution of genes within each development level obtained from Pharos[1]. **Cancer studies*:*** List and description of individual cancer studies from which our analyses are based. **Cancer Mutations in each TDL:** The number of cancers with mutations in each target development level (TDL) class related to Figure 1b. **Specific Gene Mutations*:*** Frequency of dark and light gene mutations across all cancer types for each gene. **Percent Dark Gene Mutations:** Percentage of dark gene mutations in each cancer type for each gene. **Percent Light Gene Mutations:** Percentage of light gene mutations in each cancer type for each gene. **Common Essential Genes:** Publication, antibody, and monoclonal antibody counts for common essential genes (both dark and light genes). **Non-Essential Genes:** Publication, antibody, and monoclonal antibody counts for non-essential genes (both dark and light genes).

**Supplementary File 2:** Correlation scores between mRNA expression and gene dependency scores, related to Figure 4.

**Supplementary File 3:** Dose-response of cancer cell lines: The spreadsheet contains the following results/datasets according to the sheet name. **Between Cell line Dose Responses:** mean difference comparison of the dose-responses to pathway inhibitors between the cancer cell lines that have a higher dependence on dark and light genes and those with a lower dependence on dark and light genes as defined using the CRISPR-derived gene dependence scores (see methods section). **Mutations Gene Drug Response:** mean dose-response comparison between cell lines that have a mutation(s) in a particular gene versus those that do not have a mutation(s) in that particular gene. **Pathways:** List of pathways from which our analyses are based **Pathway Inhibitors:** List of pathway inhibitors from which our analyses are based. The rest of the sheets contain drug sensitive and resistant genes for all 24 pathways (e.g., PI3K MTOR), **PI3K MTOR CRISPR:** genes that are associated with significantly increased sensitivity to different pathway inhibitors in cell lines demonstrating higher dependence on the specific gene(s) for their fitness. And genes that are associated with a significant resistance to different pathway inhibitors in cell lines demonstrating higher dependence on the specific gene(s) for their fitness. **PI3K MTOR Mutation:** genes that are associated with significantly increased sensitivity to different pathway inhibitors in cell lines that have mutations in the specific gene. And genes that are associated with significant resistance to different pathway inhibitors in cell lines that have mutations in the specific gene.

**Supplementary File 4:** Survival analysis based on mRNA expression levels of dark genes in cancer. The spreadsheet contains the following results/datasets according to the sheet name. **OS-mRNA Across Cancer Types:** Overall survival analysis between patients tumours with high and low expression of a particular dark gene calculated using the Log-rank test[2]. **DFS-mRNA Across Cancer Types:** Disease free survival analysis between patients tumours with high and low expression of a particular dark gene calculated using the Log-rank test[2]. The rest of the sheets contain overall survival analysis on the association between the mRNA expression levels of specific dark genes in each cancer type.

### References


1. Oprea, T. I. *et al.* Unexplored therapeutic opportunities in the human genome. *Nature Reviews Drug Discovery* vol. 17 317–332 Preprint at https://doi.org/10.1038/nrd.2018.14 (2018).
2. Goel, M., Kishore, J. & Khanna, P. Understanding survival analysis: Kaplan-Meier estimate. *Int J Ayurveda Res* **1**, 274 (2010).
