## Supplementary Files for "Shedding Light on the Dark Genome: Insights into the Genetic, CRISPR-based, and Pharmacological Dependencies of Human Cancers and Disease Aggressiveness"

### Supplementary Figures

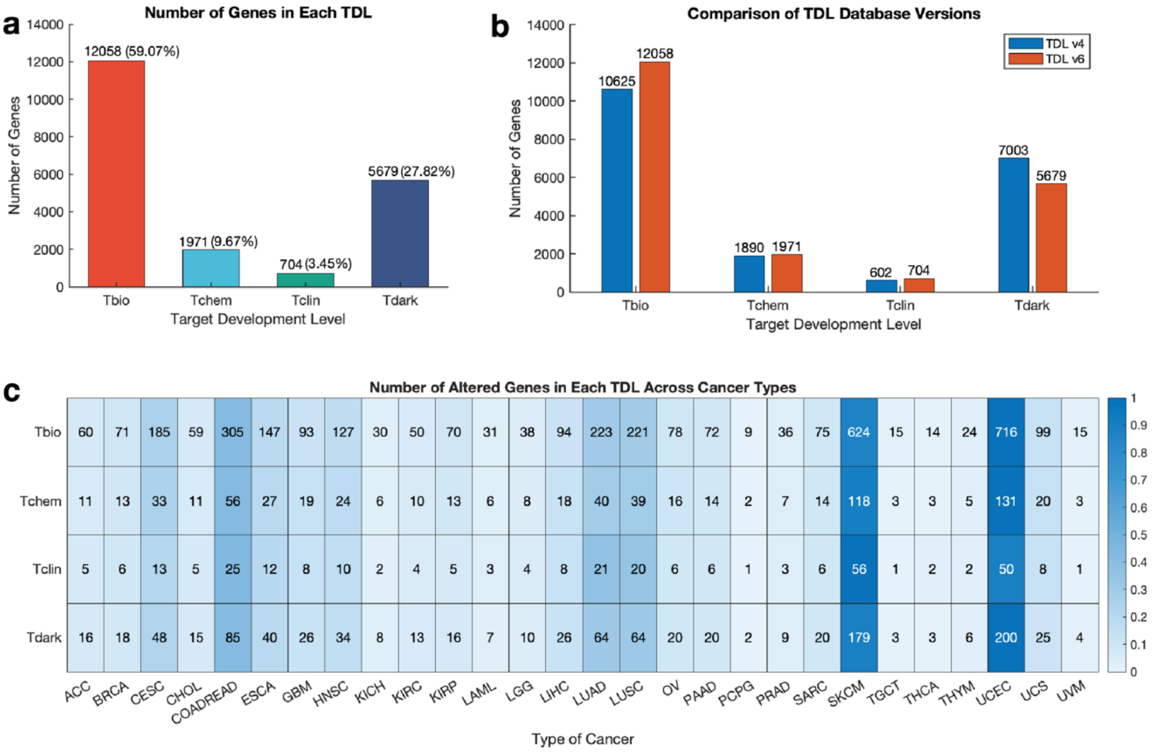


**Figure 1. Distribution of dark and light genes and their mutations in human cancers. a.** Number of genes in each target development level. **b.** Number of genes in each target development level between Target Central Resource Database (TCRD) versions 4.3.4 and 6.13.4. **c.** Number of mutated genes at each target development level across 28 cancer types. ACC: Adenoid cystic carcinoma; BRCA: Breast cancer; CESC: Cervical squamous cell carcinoma; CHOL: Cholangiocarcinoma; COADREAD: Colorectal cancer; ESCA: Oesophageal carcinoma; GBM: Glioblastoma multiforme; HNSC: Head and neck squamous cell carcinoma; KICH: Kidney chromophobe; KIRC: Kidney renal clear cell carcinoma; KIRP: Kidney renal papillary cell carcinoma; LAML: Acute myeloid leukaemia; LGG: Brain lower grade glioma; LIHC: Liver hepatocellular carcinoma; LUAD: Lung adenocarcinoma; LUSC: Lung squamous cell carcinoma; OV: Ovarian serous cystadenocarcinoma; PAAD: Pancreatic adenocarcinoma; PCPG: Pheochromocytoma and paraganglioma; PRAD: Prostate adenocarcinoma; SARC: Sarcoma; SKCM: Skin cutaneous melanoma; TGCT: Testicular germ cell tumours; THCA: Thyroid carcinoma; THYM: Thymoma; UCEC: Uterine corpus endometrial carcinoma; UCS: Uterine carcinosarcoma; UVM: Uveal melanoma.

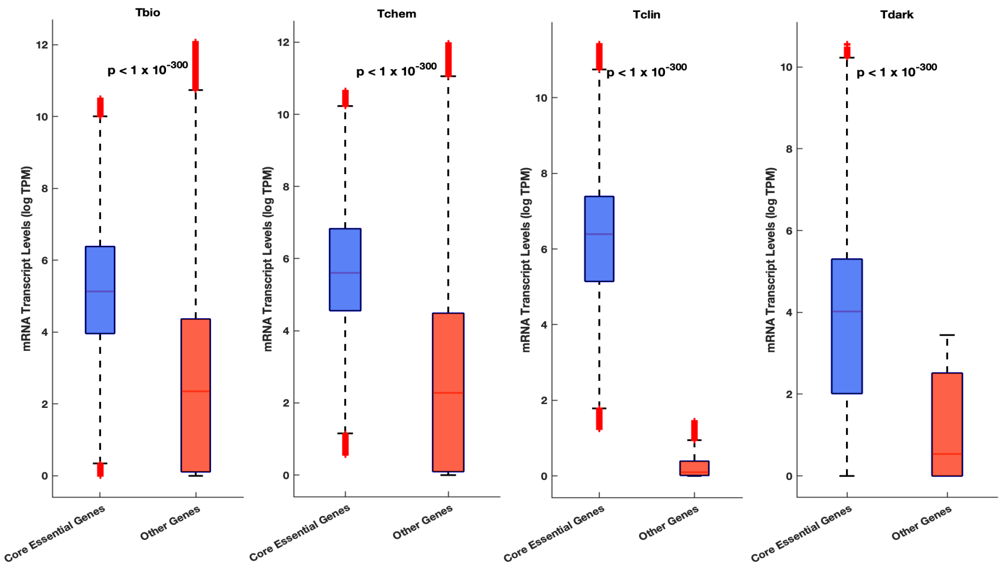


**Figure 2.** The expression of essential genes versus non-essential genes at each target development level.

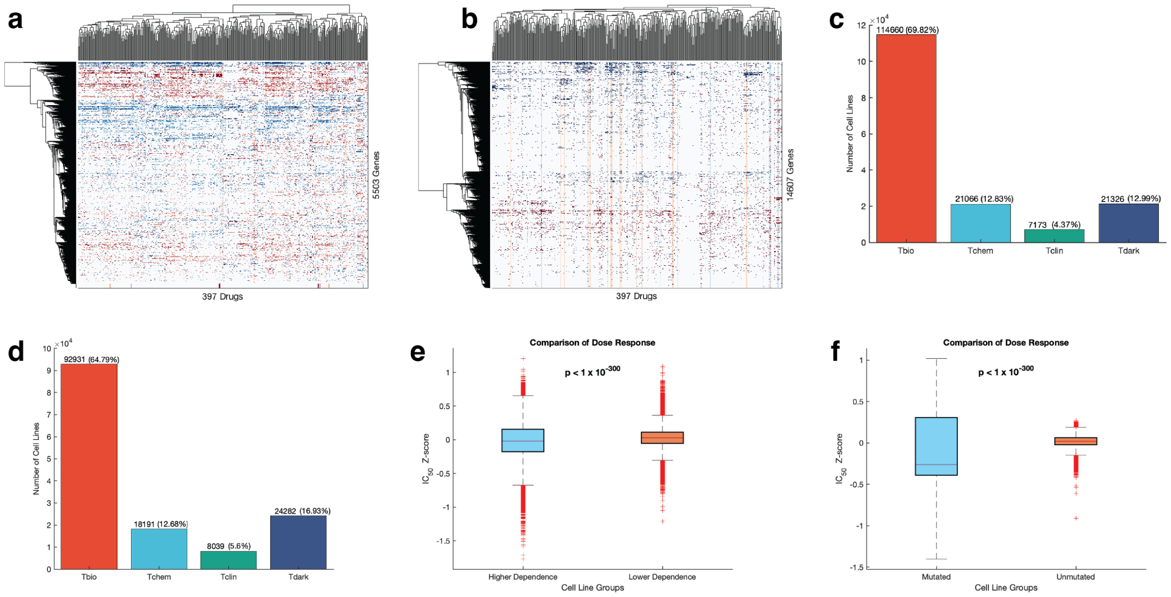


**Figure 3. Dose response of cancer cell lines. a.** The correlation between pathway inhibitors and CRISPR-derived gene dependency of cancer cell lines: analysis of 397 pathway inhibitors and 5,503 genes. **b.** The correlation between pathway inhibitors and cancer cell lines with and without specific gene mutations: analysis of 397 pathway inhibitors and 14,607 genes. **c.** The number of instances in which cell lines with varying levels of dependence on the signalling pathway significantly respond to pathway inhibitors at each target development level. **d**. The number of instances in which cell lines significantly respond to pathway inhibitors in the presence or absence of mutations in the signalling pathway at each target development level. **e.** Overall comparison of the dose-responses to pathway inhibitors between the cell lines with higher dependence on the signalling pathway and those with lower dependence on the signalling pathway. **f.** Overall comparison of the dose-responses to pathway inhibitors between the cell lines with mutations and without mutations in the signalling pathway. Boxplots show the logarithm transformed mean IC50 values of the cancer cell lines of each group. On each box, the central mark indicates the median, and the bottom edge represents the 25th percentile, whereas the top edge of the box represents 75th percentile. The whiskers extend to the most extreme data points not considered outliers, and the outliers are plotted individually using the '+' symbol.

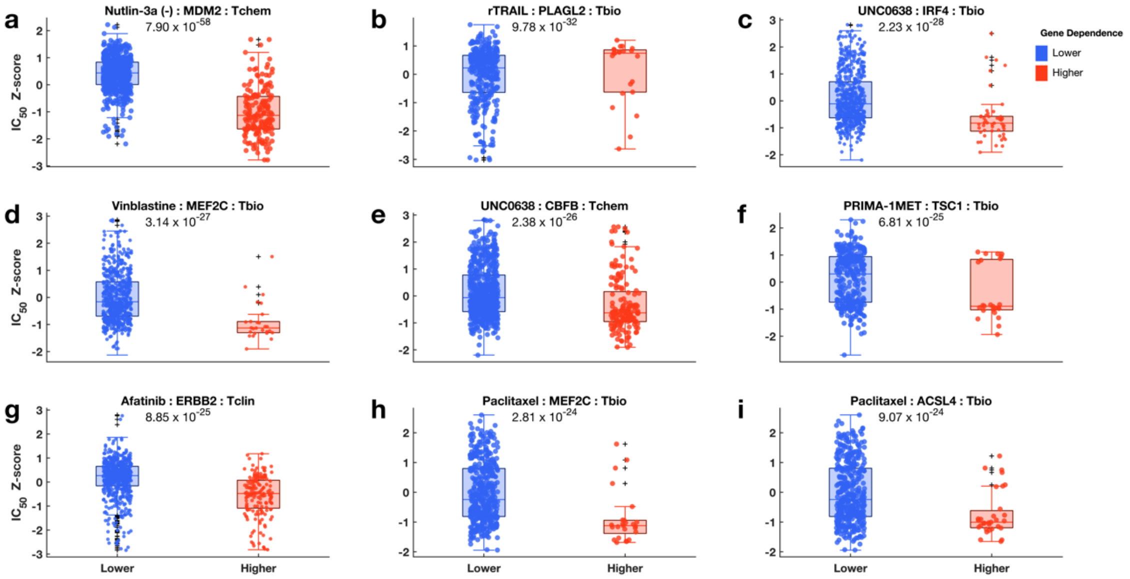


**Figure 4. Relationship between Achilles gene dependence scores and the responses of the cell lines to pathway inhibitors for light genes**. Comparison of the dose-response profiles to pathway inhibitors (**a**: Nutlin-3a (-), **b**: rTRAIL, **c**: UNC0638, **d**: Vinblastine, **e**: UNC0638, **f**: PRIMA-1MET, **g**: Afatinib, **h**: Paclitaxel, **i**: Paclitaxel) between the cancer cell lines with lower dependence (boxplots coloured blue) on signalling pathway and those with higher dependence (boxplots coloured red) on signalling pathway. Boxplots show the logarithm-transformed mean IC50 values of the cancer cell lines of each group. On each box, the central mark indicates the median, and the bottom edge represents the 25th percentile, whereas the top edge of the box represents 75th percentile. The whiskers extend to the most extreme data points not considered outliers, and the outliers are plotted individually using the '+' symbol. The scatter points within each box plot shows the overall distribution of the data points.

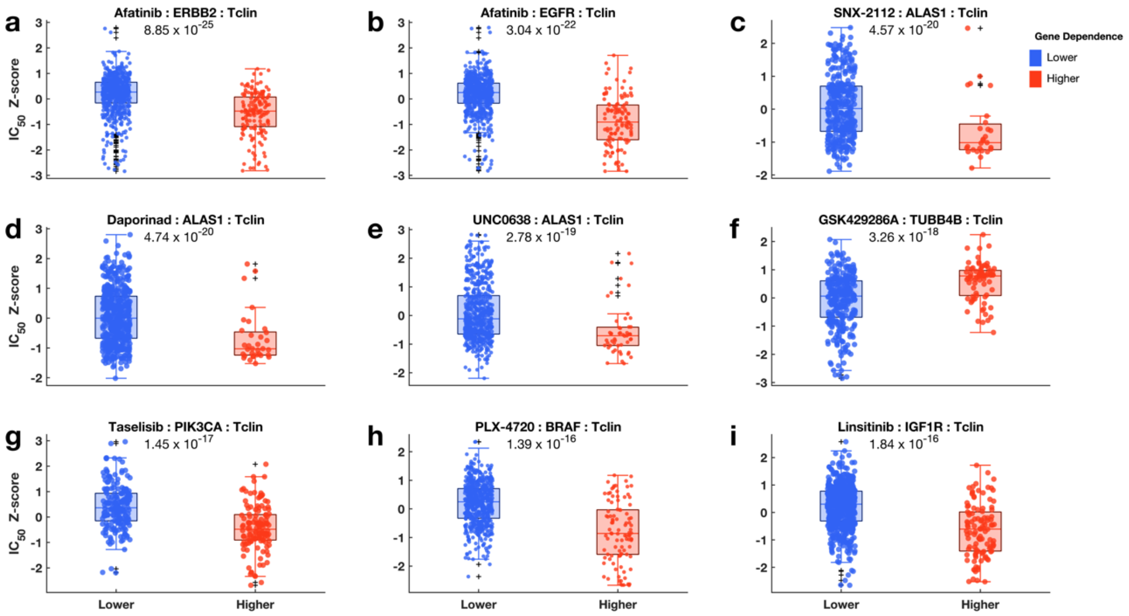


**Figure 5. Relationship between Achilles gene dependence scores and the responses of the cell lines to pathway inhibitors for Tclin genes**. Comparison of the dose-response profiles to pathway inhibitors (**a**: Afatinib, **b**: Afatinib, **c**: SNX-2112, **d**: Daporinad, **e**: UNC0638, **f**: GSK429286A, **g**: Taselisib, **h**: PLX-4720, **i**: Linsitinib) between the cancer cell lines with lower dependence (boxplots coloured blue) on signalling pathway and those with higher dependence (boxplots coloured red) on signalling pathway. Boxplots show the logarithm-transformed mean IC50 values of the cancer cell lines of each group. On each box, the central mark indicates the median, and the bottom edge represents the 25th percentile, whereas the top edge of the box represents 75th percentile. The whiskers extend to the most extreme data points not considered outliers, and the outliers are plotted individually using the '+' symbol. The scatter points within each box plot shows the overall distribution of the data points.

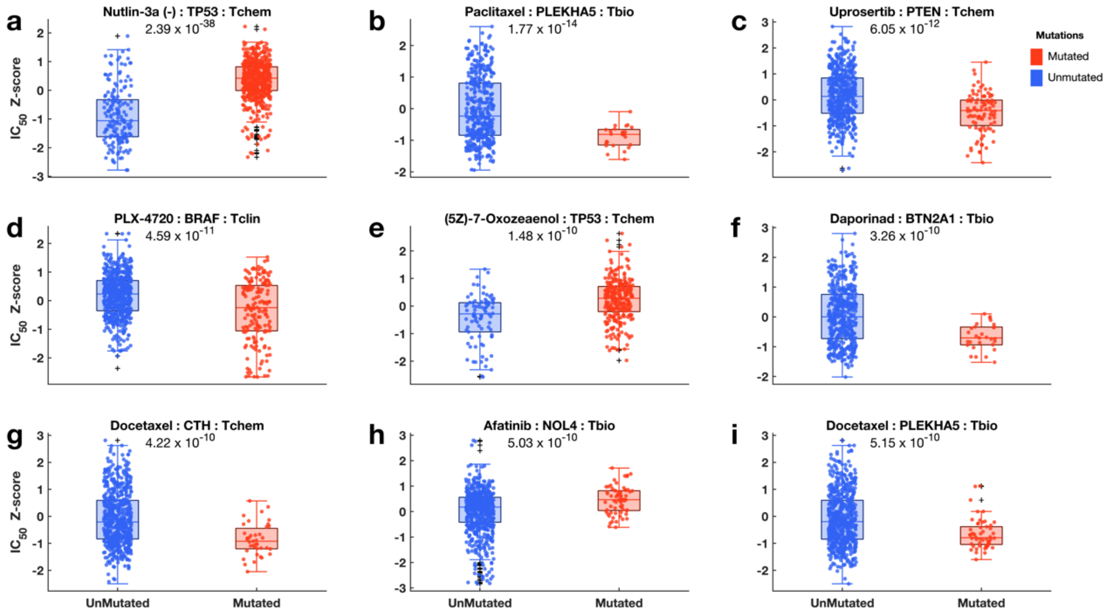


**Figure 6. Relationship between mutations of different pathway genes and the responses of the cell lines to pathway inhibitors for light genes.** Comparison of the dose-response profiles to pathway inhibitors (**a**: Nutlin-3a (-), **b**: Paclitaxel, **c**: Uprosertib, **d**: PLX-4720, **e**: (5Z)-7-Oxozeaenol, **f**: Daporinad, **g**: Docetaxel, **h**: Afatinib, **i**: Docetaxel) between the cancer cell lines without mutation (boxplots blue) on signalling pathway and those with mutation (boxplots coloured red) on signalling pathway. Boxplots show the logarithm transformed mean IC50 values of the cancer cell lines of each group. On each box, the central mark indicates the median, and the bottom edge represents the 25th percentile, whereas the top edge of the box represents 75th percentile. The whiskers extend to the most extreme data points not considered outliers, and the outliers are plotted individually using the '+' symbol. The scatter points within each box plot shows the overall distribution of the data points.

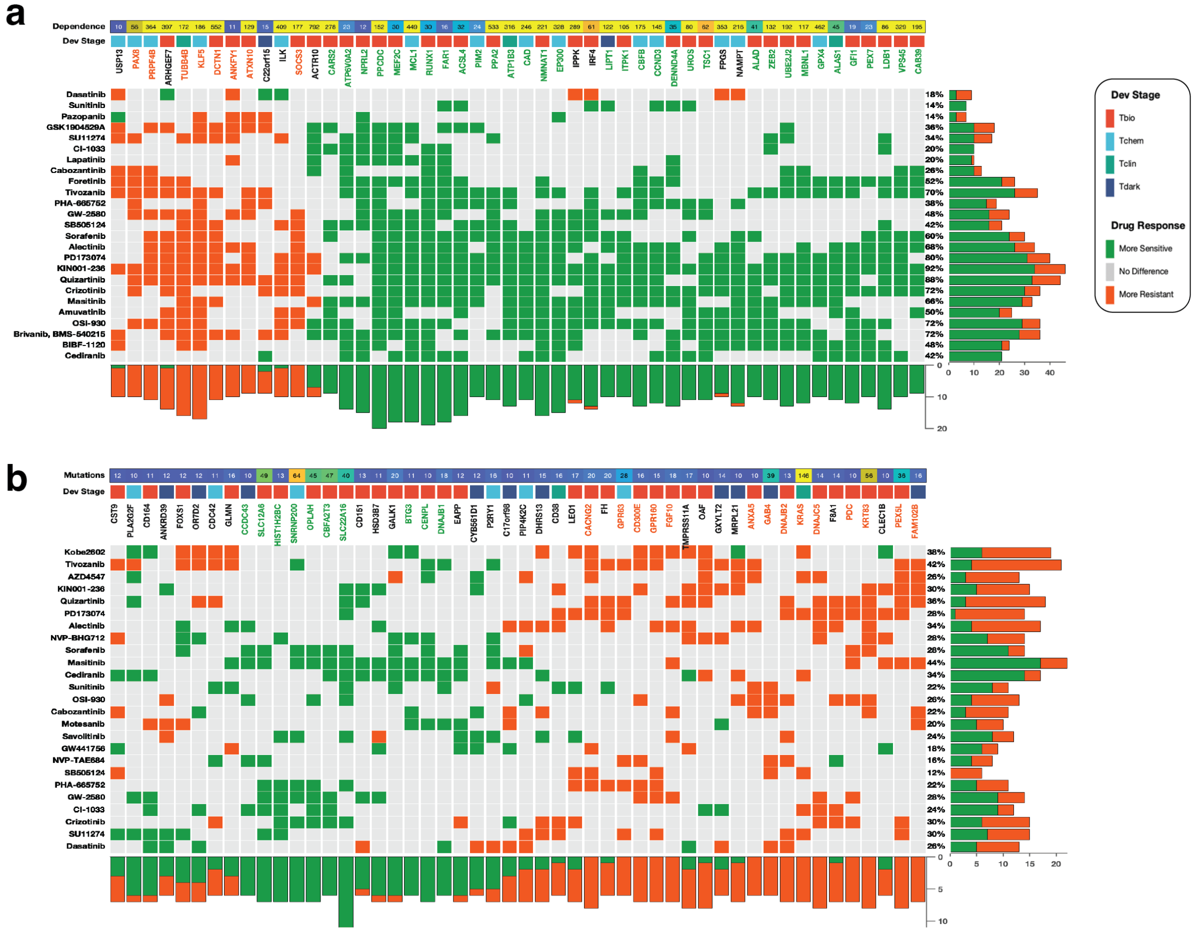


**Figure 7. RTK pathway.** The relationship between gene dependencies (**a**) or mutations (**b**) and drug responses across cancer cell lines in the RTK pathway. From top to bottom, panels indicate: Dependence; the overall CRISPR-derived gene dependence scores of the gene along that column (**a**). Overall mutation frequencies observed for the gene along the columns (**b**). Clustered heatmap; The marks on the heatmap are coloured based on how a high dependence on, or mutations in, the gene along each column affect the efficacy of the drug given along each row: (1) with green denoting significantly (10% false discovery rate) increased sensitivity, (2) grey for no statistically significant difference between cell line with a higher and lower dependence on the gene, or cell line with mutation in a gene and (3) orange denoting significantly increased resistance (for gene dependence and for gene mutations). The gene names (column labels) are coloured based on the overall calculated effect that high dependence on the gene has on the efficacy of the drug given along rows. Green: all the cell lines are significantly more sensitive to all the pathway inhibitors, orange; all the cell lines are significantly more resistant to all the pathway inhibitors, and black; a mixed response to pathway inhibitors. The bar graphs represent the total number of drugs whose dose-response is significantly increased (green) or decreased (orange).

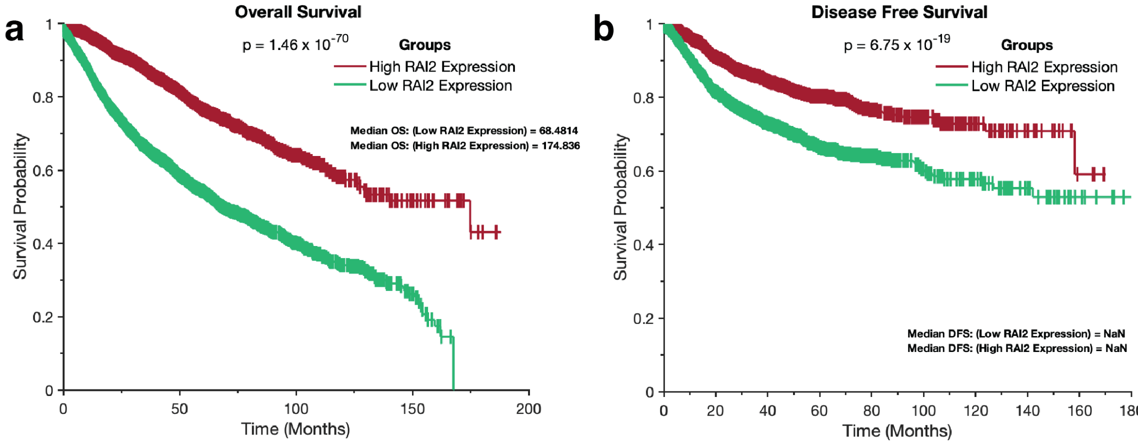


**Figure 8. Kaplan-Meir survival curves depicting the impact of dark genes on the survival of patients with cancer**. Overall survival periods (**a**) and disease-free survival periods (**b**) of TCGA patients with tumours that expressed high and low *RAI2* transcript levels.

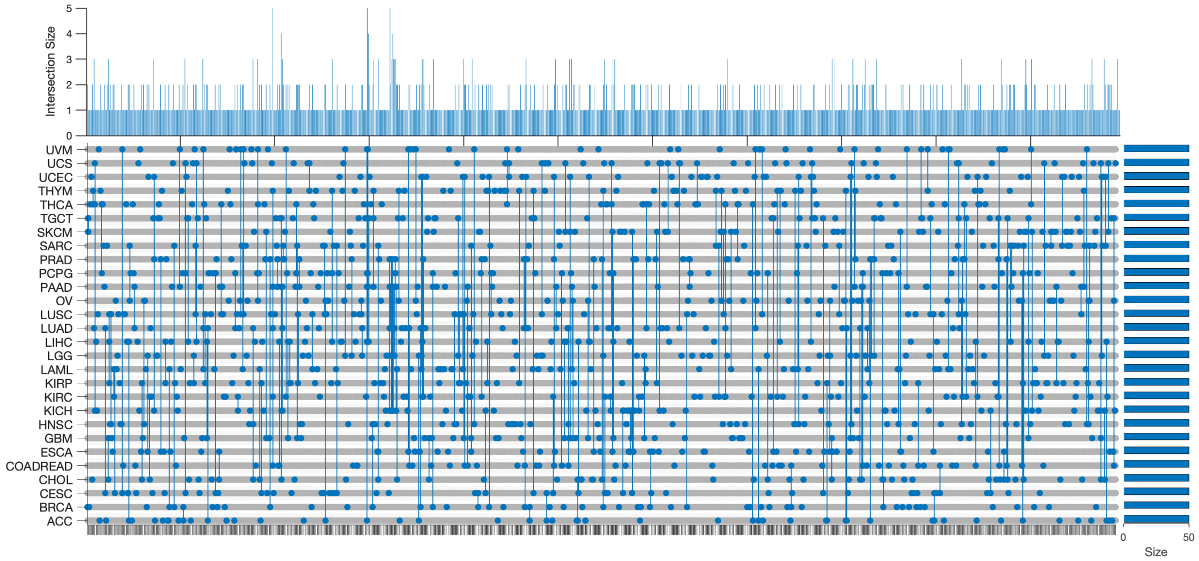


**Figure 9. UpSet plot demonstrating the distribution of the top 50 significant genes across 28 cancer types that are associated with reduced overall survival (OS), based on mRNA expression.** The plot highlights the presence of multiple overlapping genes in the screened dataset. ACC: Adenoid cystic carcinoma; BRCA: Breast cancer; CESC: Cervical squamous cell carcinoma; CHOL: Cholangiocarcinoma; COADREAD: Colorectal cancer; ESCA: Oesophageal carcinoma; GBM: Glioblastoma multiforme; HNSC: Head and neck squamous cell carcinoma; KICH: Kidney chromophobe; KIRC: Kidney renal clear cell carcinoma; KIRP: Kidney renal papillary cell carcinoma; LAML: Acute myeloid leukaemia; LGG: Brain lower grade glioma; LIHC: Liver hepatocellular carcinoma; LUAD: Lung adenocarcinoma; LUSC: Lung squamous cell carcinoma; OV: Ovarian serous cystadenocarcinoma; PAAD: Pancreatic adenocarcinoma; PCPG: Pheochromocytoma and paraganglioma; PRAD: Prostate adenocarcinoma; SARC: Sarcoma; SKCM: Skin cutaneous melanoma; TGCT: Testicular germ cell tumours; THCA: Thyroid carcinoma; THYM: Thymoma; UCEC: Uterine corpus endometrial carcinoma; UCS: Uterine carcinosarcoma; UVM: Uveal melanoma.

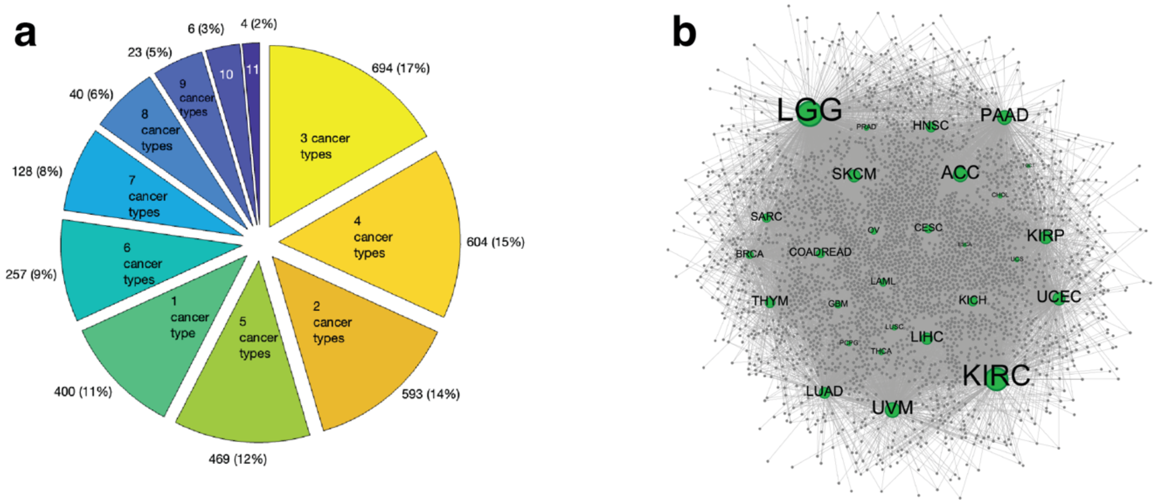


**Figure 10. Distribution of genes and their interactions with cancer types. a.** Number of significant genes with mRNA expression associated with reduced overall survival (OS) in different cancer types. **b.** Gene-cancer type interaction network. Cancer types are coloured green and genes grey. ACC: Adenoid cystic carcinoma; BRCA: Breast cancer; CESC: Cervical squamous cell carcinoma; CHOL: Cholangiocarcinoma; COADREAD: Colorectal cancer; ESCA: Oesophageal carcinoma; GBM: Glioblastoma multiforme; HNSC: Head and neck squamous cell carcinoma; KICH: Kidney chromophobe; KIRC: Kidney renal clear cell carcinoma; KIRP: Kidney renal papillary cell carcinoma; LAML: Acute myeloid leukaemia; LGG: Brain lower grade glioma; LIHC: Liver hepatocellular carcinoma; LUAD: Lung adenocarcinoma; LUSC: Lung squamous cell carcinoma; OV: Ovarian serous cystadenocarcinoma; PAAD: Pancreatic adenocarcinoma; PCPG: Pheochromocytoma and paraganglioma; PRAD: Prostate adenocarcinoma; SARC: Sarcoma; SKCM: Skin cutaneous melanoma; TGCT: Testicular germ cell tumours; THCA: Thyroid carcinoma; THYM: Thymoma; UCEC: Uterine corpus endometrial carcinoma; UCS: Uterine carcinosarcoma; UVM: Uveal melanoma.
